# Co-Contraction Embodies Uncertainty: An Optimal Feedforward Strategy for Robust Motor Control

**DOI:** 10.1101/2024.06.17.599269

**Authors:** Bastien Berret, Dorian Verdel, Etienne Burdet, Frédéric Jean

**Affiliations:** CIAMS, Université Paris-Saclay, F-91405 Orsay, France; CIAMS, Université d’Orléans, F-45067 Orléans, France; Imperial College of Science, Technology and Medicine, W12 0BZ London, United-Kingdom; Unité de Mathématiques Appliquées, ENSTA Paris, Institut Polytechnique de Paris, F-91120 Palaiseau, France

**Keywords:** optimality, co-contraction, feedforward, open-loop, stochasticity, randomness

## Abstract

Despite our environment is often uncertain, we generally manage to generate stable motor behaviors. While reactive control plays a major role in this achievement, proactive control is critical to cope with the substantial noise and delays that affect neuromusculoskeletal systems. In particular, muscle co-contraction is exploited to robustify feedforward motor commands against internal sensorimotor noise as was revealed by stochastic optimal open-loop control modeling. Here, we extend this framework to neuromusculoskeletal systems subjected to random disturbances originating from the environment. The analytical derivation and numerical simulations predict a singular relationship between the degree of uncertainty in the task at hand and the optimal level of anticipatory co-contraction. This prediction is confirmed through a single-joint pointing task experiment where an external torque is applied to the wrist near the end of the reaching movement with varying probabilities across blocks of trials. We conclude that uncertainty calls for impedance control via proactive muscle co-contraction to stabilize behaviors when reactive control is insufficient for task success.

**Author summary:** This work presents a computational framework for predicting how humans modulate muscle co-contraction to cope with uncertainties of different origins. In our neuromusculoskeletal system, uncertainties have both internal (sensorimotor noise) and external (environmental randomness) origins. The present study focuses on the latter type of uncertainty, which had not been dealt with systematically previously despite its importance in everyday life. Therefore, we thoroughly investigated how random disturbances occurring with some probability in a motor task shape the feedforward control of mechanical impedance through muscle co-contraction. Here we provide theoretical, numerical and experimental evidence that the optimal level of co-contraction steeply increases with the uncertainty of our environment. These findings show that muscle co-contraction embodies uncertainty and optimally mitigates its consequences on task execution when feedback control is insufficient due to sensory noise and delays.

## Introduction

The co-contraction of muscles spanning a joint has long been studied in human motor control (see [1] for a review). Although metabolically costly, humans will rely on an anticipatory –possibly transient– muscle co-contraction to perform a motor task in various occasions. For instance, muscle co-contraction is increased when walking on uneven ground than on on flat ground [2]. For upper-limb movements, the use of muscle co-contraction has been well characterized in tasks involving adaptation to unstable dynamics typical of tool use [3–6]. Whether co-contraction serves to modulate the mechanical impedance of the system [3] or to make feedback control more efficient [7, 8] (for instance via scaling up gains [9, 10] or enhancing response times of muscles [11]), it does contribute to robustify motor behaviors by making them more stable, accurate and reproducible [12–14]. However, few models of neuromechanical control provide a principled account of muscle co-contraction. The classical stochastic optimal control theory does not predict well co-contraction as it would constitute a waste of energy compared to the efficient feedback control [15,16]. Yet, there are substantial noise and delays in the central nervous system (CNS) [17,18] so that pure feedback control may not be a viable strategy for the task at hand. Other models have proposed that feedforward control can generate robust motor behaviors, especially with variable impedance systems like the human neuromusculoskeletal system [19–21]. When considering the noise and delays in feedback loops, feedforward co-contraction can even constitute a minimum-effort strategy [22, 23].

If most computational models focus on the uncertainty arising from within the CNS, uncertainty can also come from the environment and trigger motor adaptations (e.g., [24–28]). For instance, when exposed to a force field that is randomly turned on or off in consecutive trials, humans tend to co-contract the muscles of the relevant joints in anticipation as a response [29,30]. Incidentally, the very estimation of mechanical impedance requires the application of unpredictable disturbances to human limbs (e.g., [31]). Since humans usually adapt to such disturbances by increasing their mechanical impedance, this illustrates how uncertainty and impedance are intricately connected quantities. Therefore, the development of computational models to predict how the CNS should modulate muscle co-contraction as a function of task uncertainty will shed light on this ubiquitous motor strategy.

Here, we extend our previous framework of stochastic optimal open-loop control [20,21,32] to handle both internal and external types of uncertainty. Importantly, this framework can be applied to the nonlinear neuromusculoskeletal system. This partly comes from the restriction to open-loop control, which allows us to derive efficient methods for computing the optimal level of feedforward co-contraction given the task uncertainty, by leveraging the tools of deterministic optimal control. Our theoretical analysis predicts the existence of a logarithmic relationship between environmental uncertainty and muscle co-contraction, so that co-contraction should steeply increase with the degree of task uncertainty. This is tested in an experiment with human participants, which confirms the plausibility of the theory.

## Results

### Uncertain stochastic optimal open-loop control

The proposed framework assumes a nonlinear control system subjected to uncertainties arising from the sensorimotor noise in the CNS and the randomness of the environment. These two sources of uncertainty differ in their nature, hence are modeled distinctly. More precisely, we shall consider the nonlinear stochastic differential equation in Itô’s sense [33]

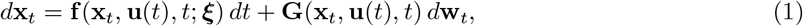

where **x**_*t*_ ∈ ℝ^*n*^ is the state vector (including position, velocity etc.), **u**(*t*) ∈ ℝ^*m*^ the control vector (e.g., muscle activations or torques), ***ξ*** a *p*-dimensional random vector modeling the environmental uncertainty (discrete or continuous with mean ***µ*** and covariance **Σ**) and **w** a multi-dimensional Wiener process modeling the internal uncertainty in the CNS [16, 34, 35]. The drift **f** and diffusion **G** are smooth nonlinear functions. In this framework, each trajectory **x**_*t*_ generated by the control **u**(*t*) depends on the fluctuations of the random variable ***ξ***, the Wiener process **w** and the random initial state **x**_0_ (with mean **m**_0_ and covariance **P**_0_, which could originate from a state estimation procedure not modeled here). Throughout the paper, we will use the notation **x**_*t*_ to distinguish the stochastic process solution to a stochastic differential equations (SDE) from the random time function **x**(*t*) solution to an ordinary differential equations (ODE). The latter occurs when the diffusion term **G** is null. Furthermore, since we focus on the role of feedforward motor commands, we explicitly assume that the control is open-loop throughout the paper, and writes it as the deterministic function **u**(*t*) for *t* ∈ [0, *T* ] where *T* is the length of the time horizon.

We assume that the motor planning process aims at finding the open-loop control **u** that minimizes the cost function

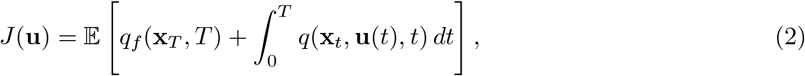

where *q* and *q*_*f*_ are quadratic functions in the state **x** and 𝔼 [*·*] denotes the expectation with respect to the random variable (***ξ*, x**_0_, **w**).

Let us define the mean **m**(*t*) = 𝔼 [**x** ] and covariance **P**(*t*) = 𝔼 [(**x** − **m**(*t*))(**x** − **m**(*t*))^⊤^] of **x**, as well as the cross-covariance **D**(*t*) = 𝔼 [(**x**_*t*_− **m**(*t*))(***ξ*** − ***µ***)^⊤^] between **x**_*t*_ and ***ξ***. In general the random variables **x**_0_ and ***ξ*** are assumed to be uncorrelated so that **D**(0) = **0**. As shown in the Materials and Methods, the uncertain stochastic optimal open-loop control (USOOC) problem defined by Eqs. 1-2 can be approximated by the following deterministic optimal control problem in augmented state (**m, P, D**).

#### Problem 1.

*The problem defined by Eqs*. 1-2 *can be approximated by a deterministic optimal control problem with dynamics*

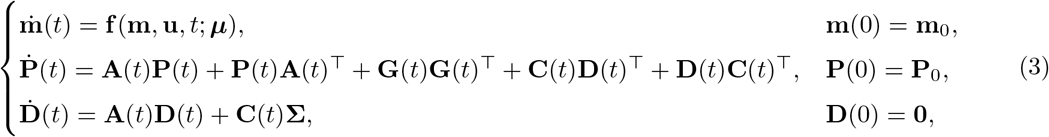

*where* **A**(*t*), **C**(*t*) *and* **G**(*t*) *are defined from Eq*. 1 as

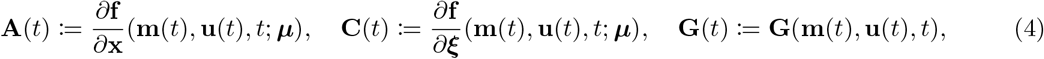

*and cost function*

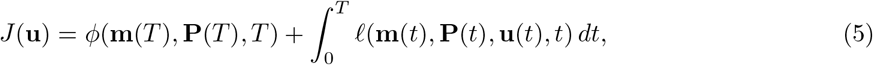

*where the functions ϕ and* 𝓁 *can be determined from Eq*. 2.

To illustrate how the cost function can be obtained, assume for instance that

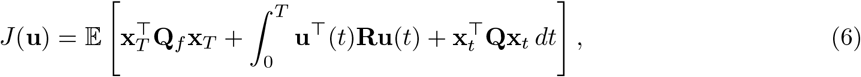

where **R** is positive definite and **Q, Q**_*f*_ are positive semidefinite matrices of appropriate dimensions. The expectation can then be rewritten only in terms of **m**(*t*) and **P**(*t*) as

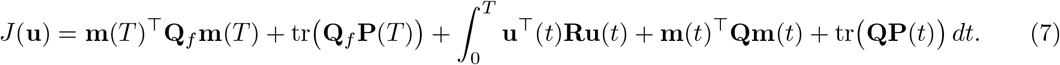

We note that a reference trajectory could be added in Eq. 6. Also, additional boundary or path constraints could be considered in more general formulations of the problem. The only requirement is to be able to write those additional constraints in terms of the mean and covariance of **x**_*t*_. For instance, a constraint on the probability to reach a given target can be added in this framework.

Importantly, the above approach yields exact solutions to the original problem when *(i)* **f** is an affine function of **x**, and *(ii)* **G** is independent on **x** (see Materials and Methods). The case where ***ξ*** is deterministic has been treated in details in [20, 21]. There, it was shown that co-contraction is an optimal strategy to minimize effort and variance objectives in presence of internal sensorimotor noise. Here, our focus is instead on the effects of the random variable ***ξ*** on the optimal feedforward motor strategy. These effects can be isolated when **G** ≡ **0** and, therefore, it is interesting to initially consider this scenario. In this case, Eq. 1 rewrites as the ODE

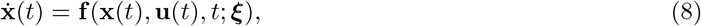

and the expectations 𝔼 [*·*] are taken with respect to the random variable (***ξ*, x**_0_). To solve this problem, the solution proposed by Problem 1 can be used by setting **G** ≡ **0**. However, an alternative approach consists of extending the original state **x** with *s* copies corresponding to different values of ***ξ***. This is readily the case for discrete variables and it can be obtained via discretization when ***ξ*** is a continuous variable (e.g., [36–41]). The dimensional advantage of our stochastic linearization approach compared to the discretization one is discussed in the Materials and Methods.

In what follows, we explore how the occurrence of random disturbances coming from the environment affect feedforward control in a variety of seminal motor control tasks, in particular the planning of mechanical impedance and muscle co-contraction.

### Case of a stabilization task

To illustrate our purpose, we first present a toy example capturing the essence of a stabilization task and allowing us to keep computations tractable analytically. Let us consider the bilinear system

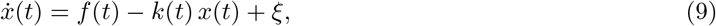

where *x* is the scalar state, **u** = (*f, k*) is the control vector composed of a term *f* representing a net “force” and a term *k* ≥ 0 representing a “stiffness”. The parameter *ξ* represents an external disturbance, which is modeled as a Bernoulli random variable with probability *α*: pr(*ξ* = 1) = *α* and pr(*ξ* = 0) = 1 − *α*. Therefore, the mean and variance of *ξ* are respectively *µ* = *α* and Σ = *α*(1 − *α*), the latter being a measure of task uncertainty. The random variable *ξ* represents a disturbance that may occur over the time horizon [0, *T* ] with probability *α* (e.g., as if an external force could randomly push our hand during repeated attempts to stabilize it).

We thus assume that the optimal control is determined as the one that minimizes the expected quadratic cost with scalar weights *q*_*f*_ *>* 0 and *q* ≥ 1,

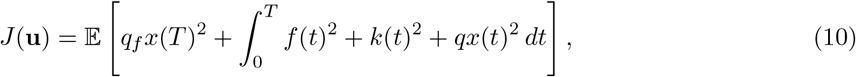

among the open-loop controls **u**(*t*) for *t* ∈ [0, *T* ] ensuring that the expectation 𝔼 [*x*(*T*)] of the final state equals 0. We assume that the initial state is *x*(0) = 0 and that time *T* is fixed.

Interestingly, it can be proven that when there is some uncertainty in the environment (i.e., 0 *< α <* 1), the optimal control verifies *k >* 0 (see Supporting Information Text for the proof). Reciprocally, when there is no uncertainty in the environment (i.e., *α* = 0 or *α* = 1), the optimal control is such that *k* ≡ 0. In conclusion, our model predicts a singular relationship between the very presence of uncertainty in the task and stiffness control.

We performed numerical simulations to visualize this relationship for varying probabilities *α* with results shown in Figure 1 (black traces). Contrary to the mean optimal net force *f* that evolves linearly depending on the probability of the external disturbance *ξ*, the evolution of the mean optimal stiffness *k* is strictly convex upwards and exhibits vertical tangents at *α* = 0 and *α* = 1. Accordingly, the evolution of stiffness with respect to task uncertainty exhibits a logarithmic shape, with a steep increase at low degrees of uncertainty (Fig. 1C). Interestingly, if we add a multiplicative noise in these simulations (10% of the input **u**), the above relationships remain valid but a lower stiffness becomes optimal. Indeed, a too large stiffness would increase uncertainty and the expected cost so that it is then better to lower the overall stiffness in this case.

**Figure 1.**
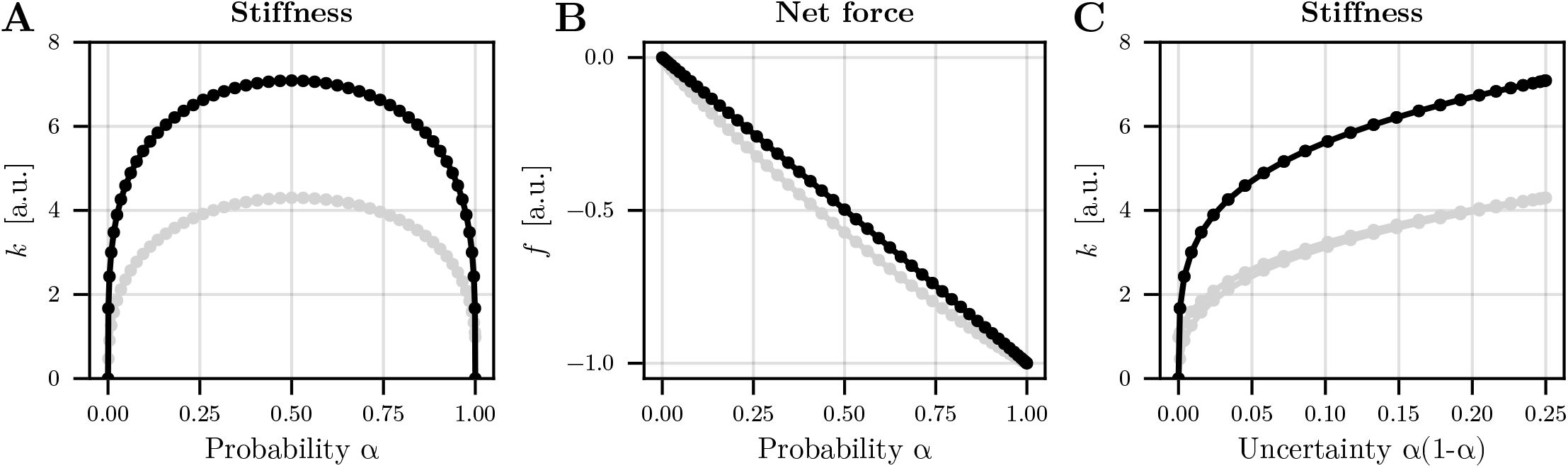
Optimal feedforward strategy for the toy stabilization task. A. Evolution of mean stiffness as a function of probability *α* which determines the occurrence of the external disturbance. Two conditions are depicted: in black, only external noise is considered; in grey, internal multiplicative noise (10%) is added. B. Evolution of mean net force as a function of probability *α*. C. Evolution of mean stiffness as a function of uncertainty, that is, the variance of the external disturbance *α*(1 − *α*). Parameters of the simulation were: *T* = 5 s, *q*_*f*_ = *q* = 10^4^. Note that the 51 values of *α* were chosen from the extrema of the Chebyshev polynomial of order 50, in order to better sample values on the edge of the [0,1] range.

To investigate further the robustness of this finding and its link with muscle co-contraction, we next simulated the inverted pendulum task studied in Hogan’s seminal work [3] but with the addition of a random external disturbance. In this task, the goal is to maintain the forearm in an upright position for 5 seconds despite the destabilizing action of gravity. In the present simulation, the system is subjected to both internal (additive motor noise) and external uncertainties (Fig. 2E). For the external uncertainty, we simulated a random disturbance taking the form of an external torque of 1 Nm applied in each trial with some probability *α* (Bernoulli variable). We varied *α* between 0 and 1. Note that the case *α* = 0 (no external disturbance) corresponds to the solution depicted in Figure 1D of the reference [21] (i.e., parameters are the same). Despite sensorimotor noise, the sharp-edged relationship between probability *α* and stiffness/co-contraction was still noticeable (Fig. 2D-E). The consideration of internal uncertainty mainly induced an offset on the latter relationship because there is a nominal level of feedforward stiffness/cocontraction even with no environmental uncertainty.

**Figure 2.**
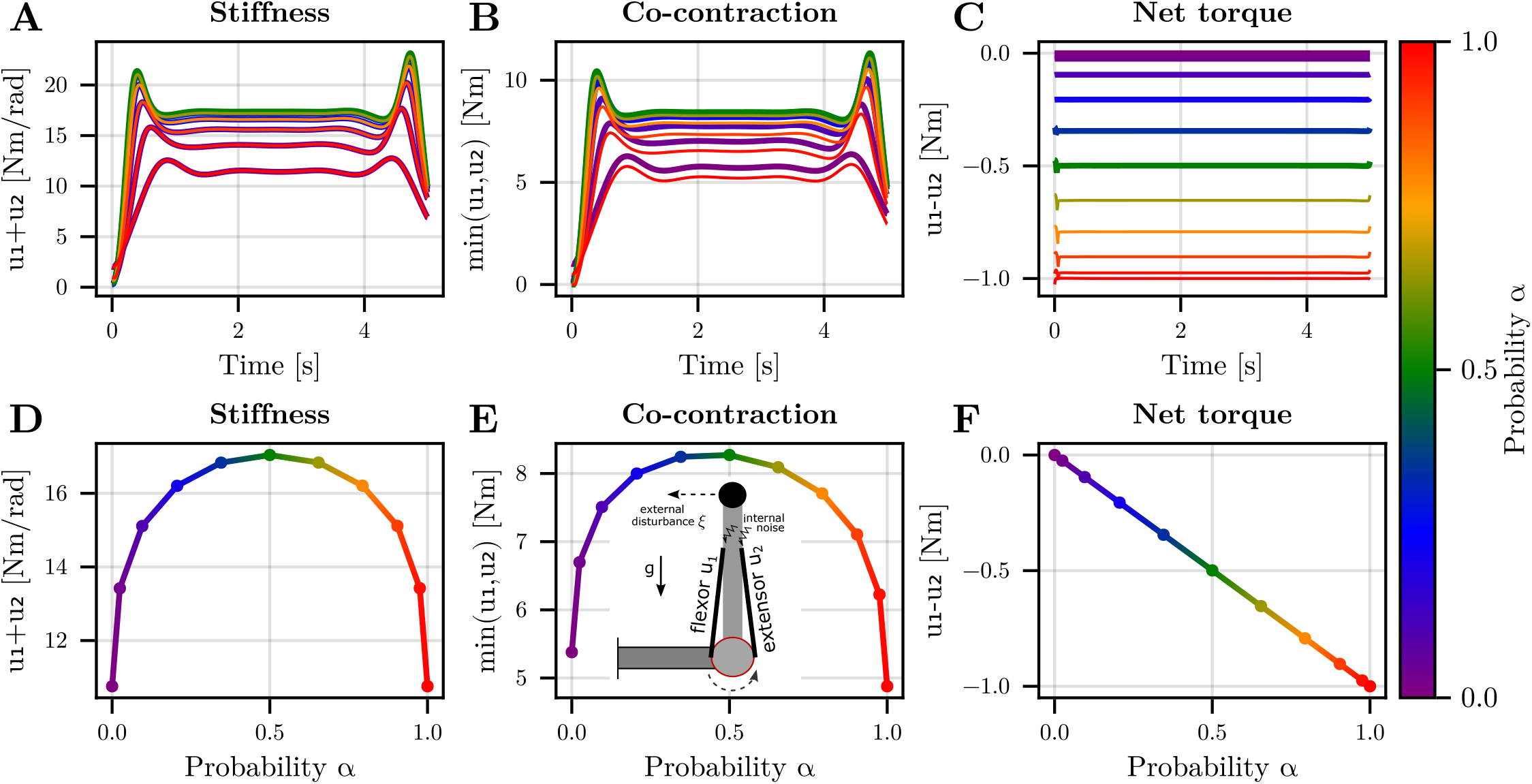
Optimal feedforward strategy for maintaining the forearm upright. A. Optimal stiffness trajectory. Stiffness is *u*_1_ + *u*_2_ where *u*_1_ and *u*_2_ are respectively the flexor and extensor elbow torques generated by muscles. The probability *α* is indicated by a color code. Note that the fluctuations surrounding the plateau are due to the initial state (small initial state covariance) and finite-time horizon (allowing specific refinements at the end of the simulation). B. Optimal level of co-contraction. Here co-contraction is simply defined as min(*u*_1_, *u*_2_). C. Optimal net torque *u*_1_ − *u*_2_. D. Evolution of mean stiffness as a function of *α*. E. Evolution of mean co-contraction as a function of *α*. The inset illustrates the task and posture. F. Evolution of mean net torque as a function of *α*. Parameters of the simulations were as in [21] (scenario with a load attached to the hand). Note that the 11 values of *α* were chosen from the extrema of the Chebyshev polynomial of order 10.

### Case of a reaching task

The two above examples consisted of simple stabilization tasks. Interestingly, as they involved bilinear drifts and quadratic cost functions, the proposed method yielded an exact solution. We show here that a similar result hold for more complex musculoskeletal systems during reaching tasks (Fig. 4). It is known that when a force field is intermittently applied across trials, participants tend to co-contract (e.g., [30]). Actually, our model suggests that the optimal level of co-contraction should change with the probability of occurrence of the force field (*α* parameter). To get some insights into how the co-contraction would vary, we simulated a planar reaching task with a two-link arm model actuated by six muscles as in the Figure 4 of reference [21]. The underlying musculoskeletal model is taken from [42]. Here, the task is to reach forward to a target (at a distance of 25 cm in 750 ms) while minimizing the control effort, Cartesian acceleration and endpoint variance, subjected to both internal and external uncertainties. The external random force field was a velocity-dependent lateral force field (i.e., a force pushing to the right with a magnitude scaling with the forward velocity of the hand) with probability *α*. Figure 3 shows that when *α* = 0 (no force field) a nominal level of stiffness and co-contraction is optimal to achieve the task. This means that the intrinsic muscle viscoelasticity is sufficient to deal with internal motor noise affecting task performance. When *α* = 0.2, that is when the force field is present in one-fifth of the trials, the optimal solution is to increase muscle co-activations, which in turn increases joint stiffness *u*_1_ + *u*_2_ and co-contraction min(*u*_1_, *u*_2_) where *u*_1_ and *u*_2_ are the normalized flexor and extensor torques. For this planar arm reaching task, the evolution of the mean stiffness and co-contraction is reported in Figure 4. It is shown that the steepness of the relationship is reduced but the pattern remains: the optimal stiffness and co-contraction must increase quite steeply as soon as some external uncertainty arises in the task before a more gentle increase is observed for larger uncertainties. This is revealed by the logarithmic curve fitting depicted in Figure 4C.

**Figure 3.**
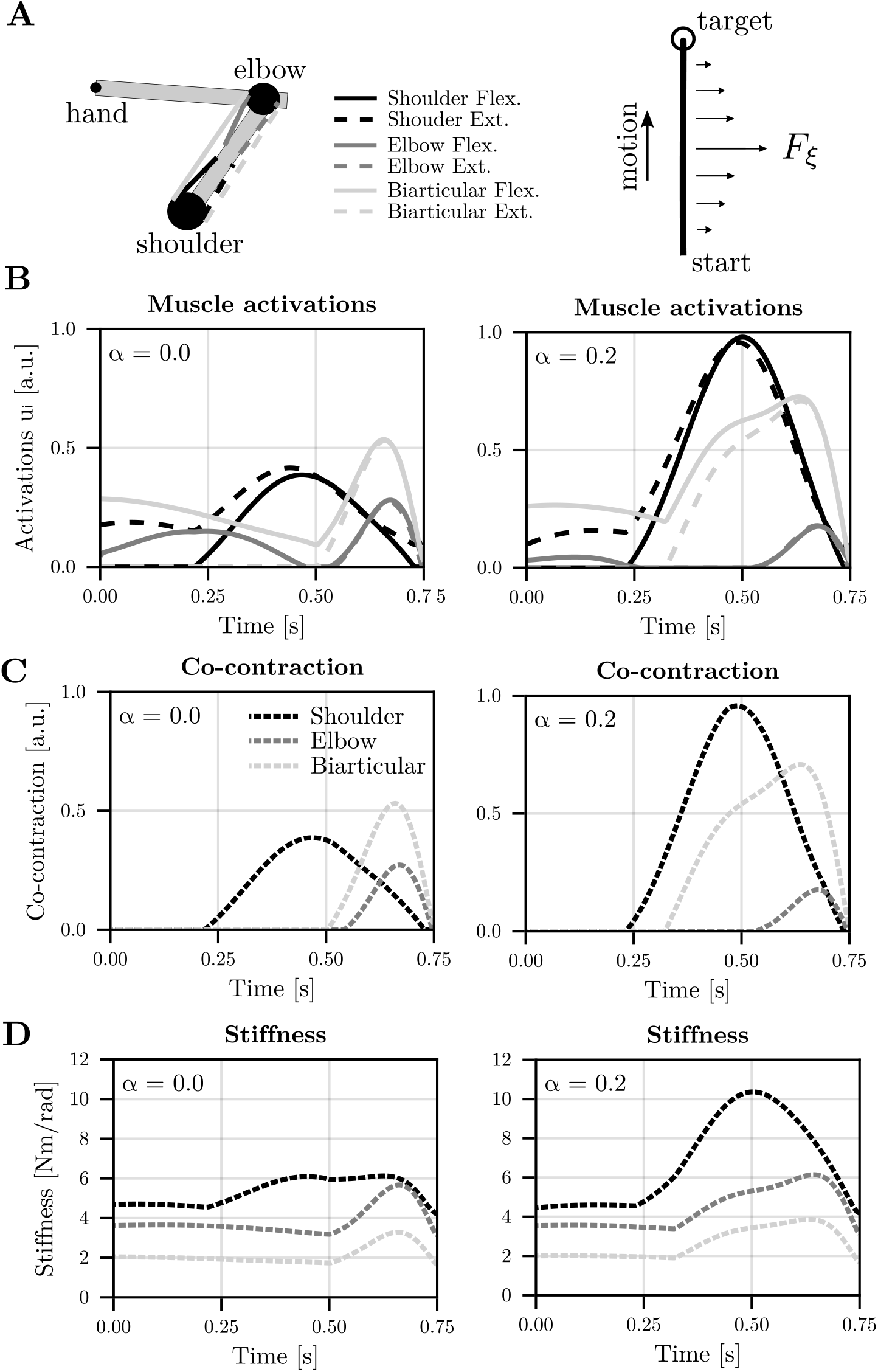
Optimal feedforward strategy for a planar arm reaching task under uncertainty. A. Illustration of the 6-muscle arm model, the initial posture and the task with the external disturbance. The horizontal force field is defined as 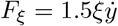 where 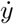 is the velocity along the y-axis of the motion. B. Optimal muscle activation pattern (for the 6 muscles) for the *α* = 0 and *α* = 0.2 conditions respectively. In the latter case, *F*_*ξ*_ is triggered with a 20% probability in each trial. C. Optimal co-contraction pattern. Each pair of muscle is treated separately (shoulder, elbow and bi-articular). D. Optimal joint stiffness pattern.

**Figure 4.**
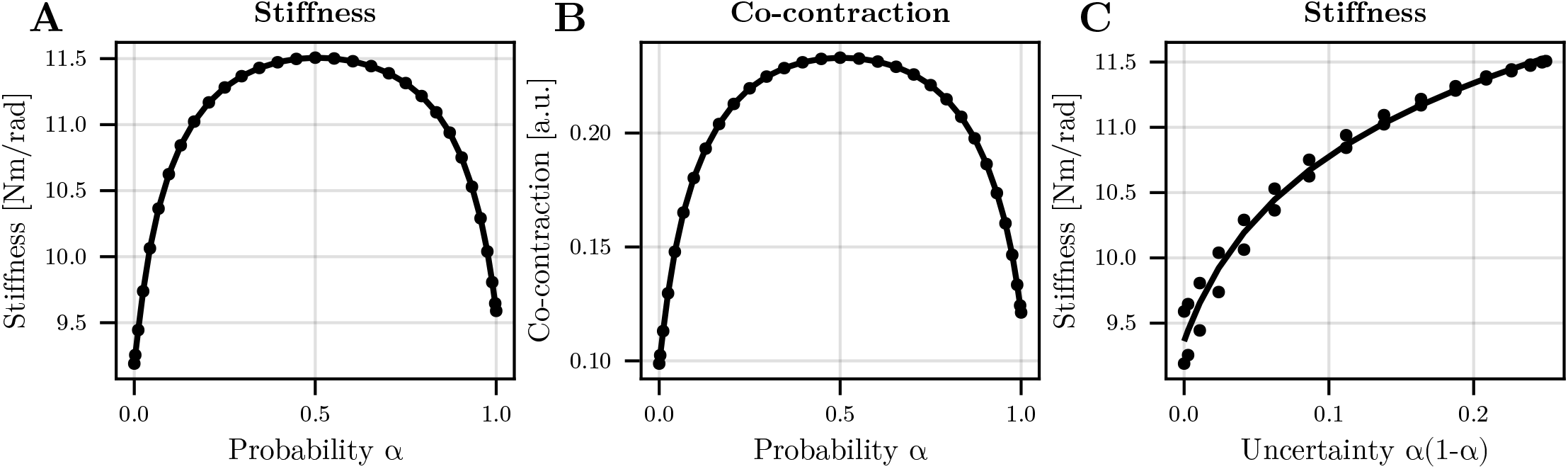
Evolution of the optimal stiffness/co-contraction. A. Evolution of mean stiffness as a function of *α* (computed as the mean stiffness from the 3 pairs of muscles). B. Evolution of mean co-contraction as a function of *α*. C. Evolution of mean stiffness as a function of external uncertainty *α*(1 − *α*). Due to the complex nonlinear dynamics, some hysteresis is observed in these simulations. A logarithm fit is used to describe the relationship between uncertainty and stiffness. Note that the 31 values of *α* were chosen from the extrema of the Chebyshev polynomial of order 30.

### Experimental testing

To test the prediction from the computational model on how co-contraction increases with random external disturbances, an experiment involving wrist flexion movements was performed with 16 participants. An active wrist exoskeleton was used to implement random disturbances with different probabilities across blocks of 100 trials (Fig. 5A-B). We designed a protocol such that the mechanical disturbances were applied near the end of the movement to emphasize the role of feedforward control. In this case, a pure feedback control strategy would fail because the neural delays are too long to ensure the target can be reached without overshooting when the disturbance suddenly occurs. Here, the disturbance was a sigmoidal torque plateauing at 0.75 Nm in 500 ms. The random disturbance was a Bernoulli variable of probability *α* ∈ *{*0, 0.25, 0.5, 0.75, 1*}*. The two first blocks without uncertainty (i.e., *α* = 0 and *α* = 1) were always performed in this order like in classical motor adaptation protocols. The three other blocks involving uncertainty were randomized across participants. Details about the experiment are given in the Materials and Methods section. As expected, participants had to adapt their motor strategy to reach the target without undershooting/overshooting in ∼500 ms as required by the protocol (Fig. 5C). Differences were observed depending on the disturbance’s probability within a block.

**Figure 5.**
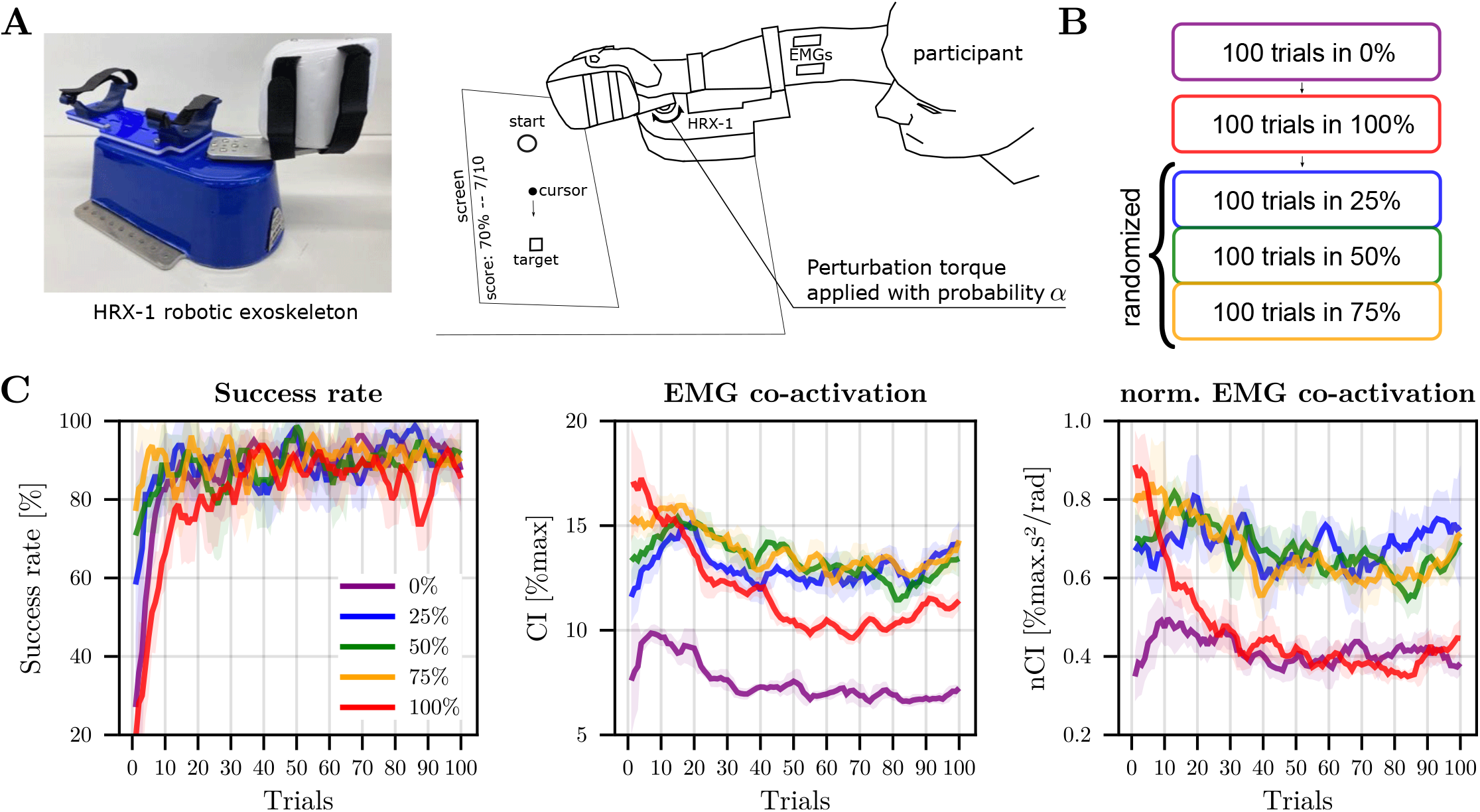
Task and behavioral adaptation across trials. A. Picture of the wrist exoskeleton robot and of the apparatus. B. Protocol with the block of trials. C. Adaptation of success rate, EMG co-activation and normalized co-activation across blocks of trials. The co-activation was computed between 150 ms before and 20 ms after the perturbation (based on the trigger position for trials without perturbation). Shaded areas represent the standard deviation across participants. The average evolution of the rate of success over 5 trials, computed over a sliding window for each participant before averaging is depicted. The corresponding trial-by-trial evolution of the muscle co-activation, averaged across participants, is also depicted as well as the normalized EMG co-activation.

The adaptation of the success rate is reported in the first panel of Figure 5C. It reveals that participants quickly learned to perform the task in absence of any perturbation (*α* = 0), with an average success rate reaching a plateau consistently above 80% after a dozen of trials. For *α* = 1, the success rate stabilized to a plateau above 80% of success rate after ∼30 trials. Interestingly, the same levels of success rates were reached for the three other conditions (i.e., *α* = 0.25, 0.5, 0.75) after only a couple of trials.

We then analyzed EMG co-activation on the window [-150;20] ms around the disturbance onset, that is, before any neural feedback could influence EMG signals. Figure 5C show that there was a clear adaptation trend in the *α* = 1 condition, with a plateau attained after ∼50 trials for the standard EMG co-activation index (CI, Eq. 25). Since EMG is known to scale with peak acceleration/deceleration [43], we also considered a normalized EMG co-activation index (nCI). Indeed, although we attempted to impose a movement time of ∼500 ms, participants tended to move slightly faster when the perturbation was present (mean movement times were: *α* = 0%: 575 ms, *α* = 25%: 574 ms, *α* = 50%: 566 ms, *α* = 75%: 562 ms, *α* = 100%: 506 ms). Thus, nCI describes more faithfully the stiffening of human joints as it removes the kinematic-dependent effects. About 30 trials were needed to attain a plateau for this normalized EMG co-activation index (nCI, Eq. 26). Interestingly, all conditions appeared to be stabilized in the second half of the block.

In order to compare adapted motor strategies, as assumed by our optimal control model, we focused on the last 50 trials with nearly constant co-activation level for the subsequent analyses. The evolution of CI and nCI as a function of the disturbance’s probability within a block are reported in Figure 6A-B.

**Figure 6.**
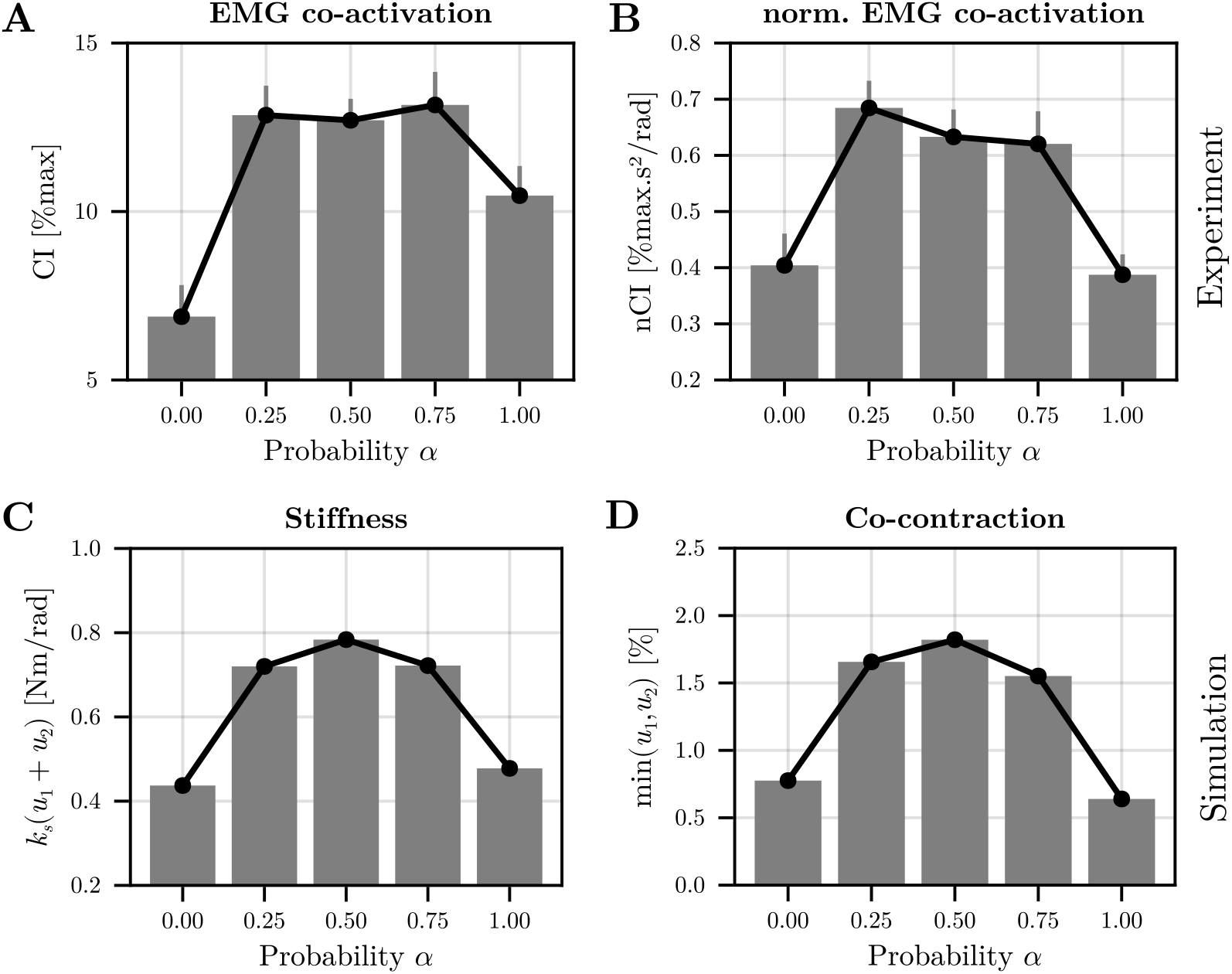
Changes in EMG co-activation as a function of the disturbance’s probability *α*. A. EMG co-activation index (CI) before any neural feedback response could influence EMG signals. Mean values across participants are reported and error bars represent the standard errors of the mean. B. Same information for the normalized EMG co-activation index (nCI). Evolution of the optimal stiffness as a function of the disturbance’s probability. C. Mean stiffness as a function of probability *α* in simulation. D. Mean co-contraction as a function of probability *α* in simulation. In simulation, the success rate was about 85%; hence similar to experimental values.

After adaptation, we observed higher values of CI when uncertainty was present in the task (0 *< α <* 1). A main effect of the disturbance’s probability on CI was found (*F*_4,60_ = 17.9, *p <* 10^−6^, *η*^2^ = 0.54). Subsequent post-hoc comparisons showed that CI values were smaller in the 0% than in all other conditions (in all cases: *p <* 0.009, Cohen’s *D >* 0.99). Furthermore, co-activation was significantly smaller in the 100% condition than in the 50% and 75% conditions (in both cases: *p <* 0.05, Cohen’s *D >* 0.72). Regarding nCI, a main effect of the disturbance’s probability was still present (*F*_4,60_ = 18.4, *p <* 10^−6^, *η*^2^ = 0.55) and nCI was significantly higher in the 25%, 50% and 75% conditions than in the 0% condition (in all cases: *p <* 0.012, Cohen’s *D >* 0.94). Importantly, these three uncertain conditions also exhibited significantly higher nCI than the 100% condition (in all cases: *p <* 0.002, Cohen’s *D >* 1.2). In summary, our results confirm that the participants significantly increased EMG co-activation when environmental uncertainty was added to the task. An important result was the significant decrease of EMG co-activation in the 100% condition compared to the 75% condition, thus showing that EMG co-activation does not simply scale with the frequency of the disturbance.

To verify if this trend was coherent with the model’s predictions, we simulated this uncertain wrist reaching task (see Materials and Methods for details). The predicted evolution of stiffness and cocontraction as a function of the probability *α* are reported in Figure 6C-D. A good match between the experimental EMG co-activation and the predicted stiffness/co-contraction can be observed, thereby showing the plausibility of theory. Because our protocol limits us to only 3 different values of the variance *α*(1 − *α*), we could not display the evolution of uncertainty as a function of stiffness using a logarithmic fit.

## Discussion

In the subway, we often hold on to the vertical bar to stabilize our body against the jolty movements of the train. When spreading our legs to widen the base of support is not possible, we can still co-contract our muscles to stiffen the arm and remain steady. The present study investigated this link between environmental uncertainty and impedance control via muscle co-contraction. To this aim, we developed a computational framework that can be applied to nonlinear musculoskeletal systems subjected to intrinsic motor noise and extrinsic random fluctuations. In particular, our modeling focused on the adaptation of the feedforward motor command to the presence of random disturbances induced by the environment. Theoretical considerations and numerical simulations led to the prediction that mechanical stiffness and muscle co-contraction steeply increase with the uncertainty in the task. To test this prediction, we conducted an experiment involving wrist reaching movements, which confirmed that EMG co-activation was greater with an uncertain force field compared to conditions where the force field was always turned on or off. Below, we discuss the implications of these findings, the limitations of our modeling and possible extensions.

We have shown that any type of uncertainty, be it internal or external to the CNS, calls for impedance control via muscle co-contraction. Our theoretical and experimental results confirmed that this is part of an optimal feedforward strategy to mitigate the effects of unpredictable disturbances. The larger muscle co-contraction when facing a greater uncertainty is coherent with a large body of the literature. First, most adaptation studies start with a baseline condition before a specific force field is suddenly turned on. Admittedly, the exposure to a novel force field can be viewed as an increase in task uncertainty. Coherently, muscle co-contraction is generally larger at the beginning of motor adaptation to a novel force field, and is progressively reduced through practice that decreases uncertainty [19, 29]. The same pattern of adaptation was also noticeable in our data for the 100% condition. Second, the study [30] found that co-contraction tends to increase when a force field is randomly turned on and off across trials, which is closely related to the type of uncertainty we considered in the present work. At last, in divergent force fields amplifying the intrinsic noise, participants exploit muscle co-contraction to adjust the endpoint stiffness, even at the end of the adaptation process [4, 6, 29]. However, in this case, it is the very presence of internal sensorimotor noise that makes such external force fields “unpredictable”. Indeed, the force field is perfectly deterministic *per se* but, for the participant, it is pushing left or right at random, hence the greater co-contraction. Indeed, simulations without sensorimotor noise do not predict any relevant co-contraction in a divergent force field (e.g., inverted forearm simulation) whereas they do in a random force field. This illustrates the conceptual difference between the uncertainty originating from the sensorimotor system itself and that from the environment.

As far as the feedforward component of the motor command is concerned, more uncertainty justifies an increase of muscle co-contraction. However, feedback is also present and crucial to human motor control as well as metabolically cheaper than co-contraction in general [15, 44, 45]. To model environmental uncertainty with feedback control, robust H-infinity control is another approach which represents a “model-free” strategy as the worst-case scenario is considered to design the control policy [30, 46, 47]. H-infinity control defines a feedback control policy to reject unmodeled disturbances, which leads to larger control gains compared to stochastic optimal control. Nevertheless, H-infinity control does not predict muscle co-contraction and has been restricted to linear systems so far, which makes it hardly applicable to more complex neuromechanical systems. In contrast, nonlinear systems can easily be considered in our framework. However, the mean and variance of the external disturbance must be estimated to optimally adjust the feedforward motor command and the level of co-contraction. This seems to be plausible with respect to the literature [25] and such an inference might be possible through adaptation, learning and inference [48, 49]. Participants may build an internal model of the task, including its uncertainty, based on their prior or recent experience [50]. Concretely here, *µ* and Σ could be meta-parameters that the brain adapts to modulate of co-contraction across repeated trials [19]. After a series of undisturbed trials, it is likely that the brain will decrease Σ whereas it will increase it following a disturbance. This could account for the trial-by-trial modulation of co-contraction found in [30]. More generally, it is likely that a combination of feedforward and feedback control will be used by the CNS to perform uncertain motor tasks [51]. Indeed, feedforward co-contraction likely provides a nominal level of stability in the task, which in turn makes feedback control more efficient by allowing larger gains and improving muscle reactivity.

It could thus be argued that the absence of feedback is a weakness in our modeling and this is a fair point. However, isolating the feedforward component of the motor command was advantageous to unveil the singular relationship between uncertainty and muscle co-contraction, and to rigorously set the foundations for future extensions. The main consequence of neglecting the effects of high-level feedback is that the predicted muscle co-contraction or stiffness is likely over-estimated in our model [52]. Indeed, sensory feedback would help to perform the task so that a lower co-contraction would probably be required in reality. In the experiment, the disturbance was actually introduced at the end of the movement so that the delays in feedback loops were too long to reject the disturbance with feedback-only control. At least two approaches can be envisioned to consider the effects of high-level feedback loops in our framework. The first one is to complement the control scheme with a locally-optimal feedback control law as in [32, 53]. While the feedback command automatically adapts to the feedforward command in this approach (e.g., the nominal stability offered by co-contraction is taken into consideration to set the feedback controller), the reverse is not true. In other words, the presence of feedback does not automatically affect the feedforward control. The second one is to consider a model predictive control approach [54–56]. Intermittent feedback control could allow the system to re-estimate its current state and plan a feedforward motor command on a receding horizon [57]. In any case, the cornerstone of high-level feedback control is the state estimation process optimally mixing sensory information and predictions with forward models [58–60]. By definition, unpredictable disturbances cannot be anticipated through forward models and, therefore, only the sensory information (innovation) provides relevant cues to update the state estimate after a random disturbance has occurred. Sensory noise and delays strongly limit the efficiency feedback control in this case. Feedforward co-contraction seems to be the solution used by the CNS to circumvent this limitation, and it fully exploits the variable impedance of muscles and the antagonistic organization of the musculoskeletal system [3]. In sum, knowing that the task is uncertain is an information that the CNS integrates to plan subsequent motor commands. Future work will focus on deriving a more comprehensive framework in which co-contraction and feedback gains are concurrently planned, taking into account the consequences of noise/delays on state estimation and feedback control with respect to the task at hand. This framework could be applied to the development of soft robot control [61] and variable impedance actuators [62], and more generally to human-robot interaction towards health and manufacturing applications [63].

## Materials and Methods

### Theoretical developments

Here, we describe the different variants of the uncertain stochastic open-loop optimal control framework, which can be used depending on the task at hand. For clarity in the presentation, we introduce the different variants by order of complexity and we start by considering the case without diffusion term **G** in Eq. 1. We shall focus first on the affine case and then on the nonlinear case.

**Affine case without diffusion** Let us consider the affine system

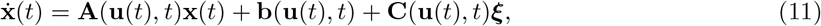

where the variables **x**(*t*) ∈ ℝ^*n*^, **u**(*t*) ∈ ℝ^*m*^, ***ξ*** ∈ ℝ^*p*^ are defined as in the Results section. The terms **A, b** and **C** are assumed to be known functions of the time *t* ∈ [0, *T* ] and of the open-loop control **u**(*t*). In practice they represent the internal model of the limb dynamics and of the task built by the CNS through learning. The cost function to minimize is in the form of Eq. 2. The problem is to find the open-loop control **u**(*t*) and the trajectory distribution **x**(*t*) starting from an initial state **x**(0) = **x**_0_ (which can be a random variable with mean **m**_0_ and covariance **P**_0_) minimizing the chosen expected cost.

To solve this problem, we will formulate an equivalent deterministic optimal control problem. To do so, let us express the ODE satisfied by the mean **m**(*t*) and covariance **P**(*t*) of **x**.

A direct computation shows that the propagation of the mean **m**(*t*) = 𝔼 [**x**(*t*)] is given by the ODE

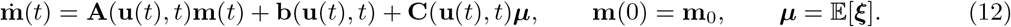

In order to get an equation for the propagation of the *n×n* covariance matrix **P**(*t*) = 𝔼[(**x**(*t*)−**m**(*t*)) (**x**(*t*)− **m**(*t*))^⊤^], we introduce the *n × p* cross-covariance matrix **D**(*t*) = 𝔼[(**x**(*t*) − **m**(*t*)) (***ξ*** − ***µ***) ^⊤^] between **x** and ***ξ***, and the *p × p* covariance **Σ** = E[(***ξ*** − ***µ***)(***ξ*** − ***µ***)^⊤^] of ***ξ***. Again, a direct computation shows that the covariance propagation is governed by the ODE

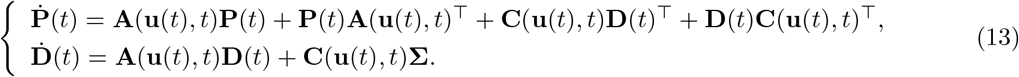

The initial conditions of **P** and **D** are respectively the covariance of the initial state **P**(0) = 𝔼[(**x**_0_ − **m**_0_)(**x**_0_ − **m**_0_) ^⊤^] = **P**_0_ and **D**(0) = 𝔼[(**x**_0_ − **m**_0_)(***ξ*** − ***µ***)^⊤^]. As already mentioned, the random variables **x**_0_ and ***ξ*** are generally uncorrelated so that we assume that **D**(0) = **0**.

#### Problem 2.

*For the affine case, the original uncertain optimal open-loop control problem can be replaced by an equivalent deterministic optimal control problem in augmented state* (**m, P, D**) *with dynamics*

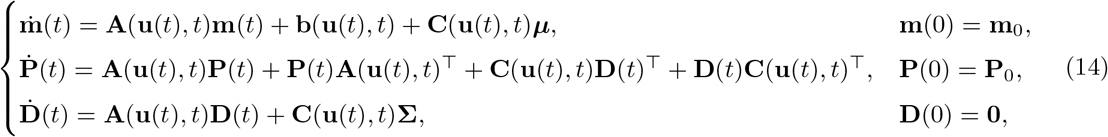

*and cost function*

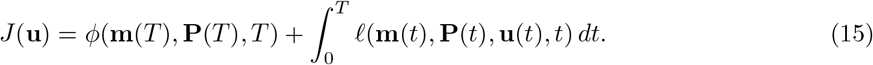

From the above, it can be noted that we made no assumption on the distribution of the random variable ***ξ***, and that the optimal solution only depends on the mean ***µ*** and covariance **Σ** of ***ξ***. Furthermore, additional boundary conditions that would only involve the mean and covariance (and possibly the cross-covariance as well) can be easily considered.

If the initial state **x**_0_ is deterministic (i.e., **x**_0_ = **m**_0_), the problem admits a much simpler expression.

Because **P**(0) = **0** and **D**(0) = **0**, we can express **P**(*t*) as a function of **D**(*t*) as follows:

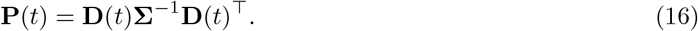

The equivalent deterministic optimal control problem drastically simplifies as it can be fully written in terms of the augmented state (**m, D**), and **P** can be directly computed from Eq. 16.

**Nonlinear case without diffusion** Let us consider the same problem but with an uncertain nonlinear system of the form

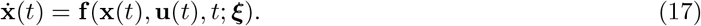

In this case, it is necessary to resort to approximation techniques. Again, we introduce the mean **m**_x_ = 𝔼[**x**] and covariance **P**_x_ = 𝔼[(**x** − **m**_x_) (**x** − **m**_x_)^⊤^] of **x**. The subscript **x** is introduced here because, in contrast to the affine case, now **m**_x_(*t*) and **P**_x_(*t*) cannot be obtained as solutions of a finite-dimensional control system. Indeed, the nonlinearity of **f** may introduce moments of any order. However, using statistical linearization techniques (see [20, 64]), we can approximate these quantities by (**m, P**)(*t*), the first two components of the trajectories (**m, P, D**)(*t*) of

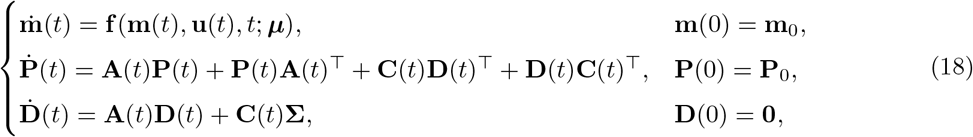

where

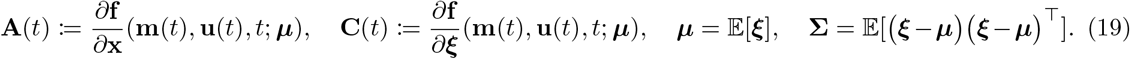

The latter system can be obtained by augmenting the system’s state with ***ξ*** and including all the uncertainty in the initial state, and then use the method in [20] with a null diffusion term (see also the proof of Lemma 1 in the Supporting Information Text). Alternatively, we can directly write the propagation of **m**_x_(*t*) and **P**_x_(*t*) as in Eq. 14 of the Supporting Information Text and use a first order Taylor series expansion to obtain the above equations.

#### Problem 3.

*For the nonlinear case, the original uncertain optimal open-loop control problem can be approximated by a deterministic optimal control problem in augmented state* (**m, P, D**) *among the trajectories* (**m, P, D**)(*t*) *satisfying Eq. 18 and minimizing the cost given in Eq. 15*.

*Remark*. In general, the dimension of the augmented state will be 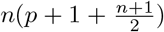. Therefore, it grows quadratically with the dimension of the state **x** but only linearly with the dimension of the random parameter ***ξ***. Interestingly, if there is no uncertainty about the initial state **x**_0_, the dimension of the augmented state reduces to *n*(*p* + 1). Alternative approaches propose to directly discretize the parameter space ***ξ*** [36–41], the expected cost being then approximated by a weighted finite sum. If *s* samples of ***ξ*** are used, an associated deterministic optimal control problem in dimension *ns* can be formulated using *s* copies of the dynamical system. We refer the reader to the Supporting Information Text where this kind of approach is explicitly developed for a discrete random variable ***ξ***. The challenge when such approaches are applied to continuous random variables is to find a small number *s* allowing accurate approximations of the expected cost. Comparing dimensions, we see that our method might be advantageous if 2*s >* 2*p*+*n*+3 or *s > p* + 1 if **x**_0_ is deterministic. If we consider the internal uncertainty coming from the diffusion term, the covariance must be added anyway in the augmented state (see Supporting Information Text), and the approach with *s* copies requires to solve a deterministic problem in 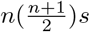 dimensions, such that our approach can become even more advantageous when *s* is too large.

In a practical setting (compactly supported dynamics), the accuracy of the approximation in Problem 3 we make is guaranteed by the following result (see Supporting Information Text for the proof and [65] for more general results of this kind).

#### Proposition 1.

*Assume that* **f** *is smooth, that the set U of control values is a compact subset of* R^*m*^, *that the random vector* ***ξ*** *takes values in a compact set, and that all the vector fields* **x** ⟼ **f** (**x, u**; ***ξ***) *have the same compact support. Let T >* 0. *Then* (**m, P**)(*·*) *converges uniformly on* [0, *T* ] *to* (**m**_x_, **P**_x_)(*·*) *when* (**P**(*·*), **D**(*·*), **Σ**) *converges uniformly to* 0.

*If moreover the dynamics is affine with respect to the random parameter* ***ξ***, *then the convergence holds independently on* **Σ**, *that is, when* (**P**(*·*), **D**(*·*)) *converges uniformly to* 0.

**Affine case with diffusion** Let us now consider the case where the dynamical system is a linear SDE with a drift that depends linearly on the uncertain parameter, that is,

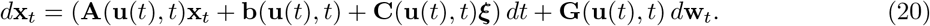

With the diffusion term, the problem is more complicated but can still be solved by formulating an equivalent deterministic optimal control problem with augmented state.

To derive the result, first remind that the cost *J*(**u**) in Eq. 2 can be rewritten as a function of the mean **m**_x_ = 𝔼 [**x**] and covariance **P**_x_ = 𝔼 [(**x** − **m**_x_) (**x** − **m**_x_)^⊤^] of the process **x**, where the expectation is taken with respect to ***ξ*, x**_0_ and **w**. Second, it is convenient to distinguish the role of the two sources of uncertainty in the following way. For a fixed instance of the parameter ***ξ***, let **x**^***ξ***^ be the solution of Eq. 1 with initial condition **x**_0_ and let 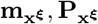 be respectively its mean and covariance with respect to (**x**_0_, **w**). Then, denoting by 𝔼_***ξ***_ the expectation with respect to ***ξ***, a computation shows that

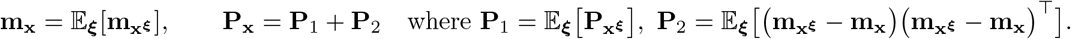

The covariance 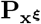 satisfies the differential equation 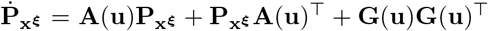 with **P**_0_ = 𝔼 [(**x**_0_ − **m**_0_)(**x**_0_ − **m**_0_ ^⊤^] as an initial condition. Neither the differential equation nor the initial condition depend on ***ξ***, so 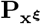 does not depend on ***ξ***, which implies that **P**_1_ is obtained as the solution of

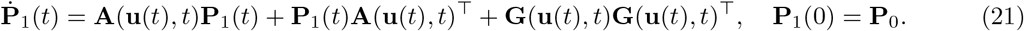

On the other hand, 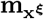 is solution of the ODE

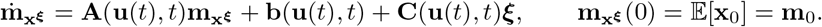

Arguing as in the affine case without diffusion, we obtain that (**m**_x_, **P**_2_) are the first two components of the trajectories (**m**_x_, **P**_2_, **D**)(*t*) of

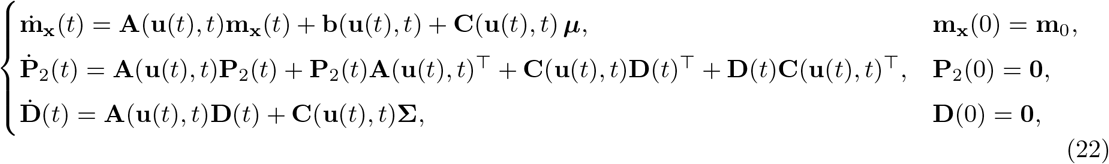

where ***µ*** and **Σ** are respectively the mean and covariance of ***ξ***.

#### Problem 4.

*For the affine case with a diffusion term, the original uncertain stochastic optimal open-loop control problem is equivalent to the following deterministic one: minimize the cost J*(**u**) *written as a function of* **m**_x_ *and* **P**_x_ = **P**_1_ + **P**_2_ *as in Eq. 15 among the trajectories* (**m**_x_, **P**_1_, **P**_2_, **D**)(*t*) *of the equations (21)-(22)*.

**Nonlinear case with diffusion** For the general nonlinear case where the system writes

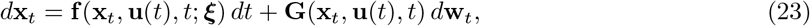

we can combine the results of [20] and of the nonlinear case without diffusion described above to propose a deterministic optimal problem approximating the original one. This deterministic problem is constructed as follows: its cost is obtained by replacing in Eq. 15 the mean **m**_x_ and covariance **P**_x_ by **m** and **P** = **P**_1_ + **P**_2_ respectively, and (**m, P, D**)(*t*) are obtained by combining Eqs. 18-21-22 to obtain the USOOC problem approximation given in Problem 1.

### Uncertain wrist reaching experiment

**Experimental protocol** A total of *N* = 16 participants were recruited for the experiment and test the model’s prediction. They were healthy right-handed adults with the following anthropometric characteristics: age 27 *±* 5 years old, height 177 *±* 7.1 cm, weight 73 *±* 8.95 kg, hand length 19.7 *±* 1 cm. The hand length was measured for each participant between the tip of the middle finger and the wrist centre (estimated as the middle of the segment formed by the head of the ulna and the radial styloid process). The protocol was approved by the Université Paris-Saclay ethical committee for research (CER-Paris-Saclay-2021-048/A1). This study used the HRX-1 (HumanRobotix, London, UK) wrist exoskeleton to record the motion and implement the mechanical disturbance. This device is a 1-degree-of-freedom active wrist exoskeleton controlled at 100 Hz. It is equipped with an encoder (Encoder MILE 512-6400, 6400 counts per turn) to measure the position of the human wrist. The participant’s hand and forearm were each attached to the exoskeleton with two cuffs. Their respective positions were adjusted closer or further from the exoskeleton’s joint axis to match the participant’s forearm length.

The task involved 60*°* wrist flexion and extension reaching movements centered around 0*°*, which corresponds to the hand aligned with the forearm. Participants were asked to reach targets that were displayed on a screen as 3-cm long green squares. Two targets were considered: left (wrist flexion posture) and right (wrist extension posture). For both targets, participants had to remain static and inside the target during 1.5 s to validate the trial. Whenever a target was validated, it disappeared and the other target appeared. The right target was always green whereas the left target was initially blue for 500 ms, then green for 500 ms and then blue again until target validation. For a successful trial, the participant had to attain the left target during the green period but could start moving during the first blue period, yielding valid movements for durations between 0.5 s and 1 s when considering both reaction time and motion time. If the target was attained before or after the green period, the trial was failed and if an overshoot was detected, the left target turned red and the trial was failed.

During the flexion movements, a mechanical disturbance was triggered at 90% of movement amplitude with a given probability in a series of trials. Five different probabilities of disturbance were tested across 5 separate blocks of 100 trials. Hence 500 flexion movements per participant were recorded and analyzed. The disturbance was presented with a Bernoulli distributed probability *α*. In the first block, no disturbance was applied (*α* = 0), so that participants could familiarize with the task and perform it seamlessly. In the second block, the disturbance was always present (*α* = 1) so that participants could adapt to it and potentially learn to compensate it. The three next blocks were randomized and the disturbance probability in these blocks was one of the following: *α* ∈ *{*0.25, 0.5, 0.75*}*. To limit muscle fatigue, 3-minute breaks were imposed between blocks. In order to motivate participants, their instantaneous rate of success during the block was displayed (both absolute and percentage) and they were told a fictitious highest success rate obtained by the “best participant” so far (selected between 90% and 95%).

The disturbance applied by the HRX-1 robot to the human wrist took the form of a sigmoidal torque with a plateau at *τ*_max_ = 0.75 Nm that was reached in roughly 500 ms. When the 90% threshold was crossed, the disturbance torque was generated as follows:

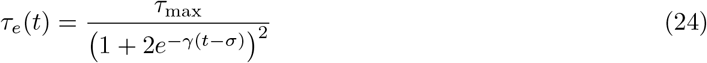

where *γ* = 9.9903 and *σ* = 0.0652. These values were chosen to ensure 500 ms between 5% and 95% of the plateau. Note that the disturbance plateau was maintained longer than the time necessary to validate a trial (i.e., 2.5 s). Thus, the release of the disturbance could not impact the performance in the task.

To assess muscle co-activations, the muscle activity of the flexor carpi radialis (wrist flexor) and of the extensor carpi radialis (wrist extensor) were recorded using bipolar surface EMG (Wave Plus, Wireless EMG, sample rate 2 kHz; Cometa, Bareggio, Italy). The EMGs were placed according to the SENIAM recommendations [66]. Before placing the electrodes, the skin was locally shaved and cleaned with a hydro-alcoholic solution.

**Data processing** EMG signals were band-passed filtered (Butterworth, 4^*th*^ order, [20; 450] Hz cut-off frequencies), centered and rectified [67]. They were then normalized by the maximum value obtained for each muscle over the course of the whole experiment. The averaged sum of the two signals over different time windows of interest were used as an EMG co-activation index, defined as

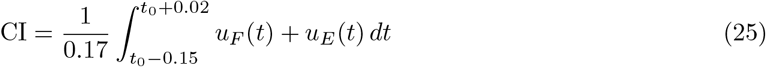

where *t*_0_ is the onset time of the disturbance, and *u*_*F*_ (*t*) and *u*_*E*_(*t*) are processed EMG signals of the flexor and extensor respectively. Note that this time window was chosen such that only anticipatory EMG activity was analyzed (hence it excludes any reflex occurring after the mechanical perturbation).

Given that muscle activity is known to correlate with joint acceleration/deceleration for single-joint movements [43], we also computed a normalized EMG co-activation index that accounts for motion deceleration as follows:

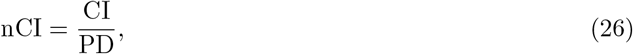

where PD is the peak of deceleration of the reaching considered movement. This normalization was chosen because the disturbance was applied near the end of the movement, that is, during the deceleration phase.

Movements were segmented based on motion kinematics as described below. Wrist joint angles were measured at 100 Hz with the encoder of the HRX-1 exoskeleton. Successive positions of the wrist were low-pass filtered (Butterworth, 5^*th*^ order, 5 Hz cut-off frequency). Wrist joint angular velocity and acceleration were obtained through numerical differentiation. Individual movements were first isolated based on the time spent by participants inside targets. Then, for each movement, initial and final times were computed using a threshold at 5% of the peak wrist angular velocity. The kinematics, task events (i.e. targets appearing and disappearing) and muscle activities were all synchronously collected using a Matlab (R2023b, Mathworks, USA) custom code.

**Statistical analyses** Main effects of the level of uncertainty were first assessed using one-way repeated measurements ANOVAs. In case sphericity conditions were not satisfied (i.e. *ϵ <* 0.75), a Greenhouse-Geisser correction was applied. For all significant ANOVA, we report the *η*^2^ as a measure of the effect size. The significance level of ANOVA was set at *p <* 0.05. In case a main effect was found, we performed pairwise *t*-tests between the different levels of uncertainty. For all significant comparisons, we report the Cohen’s D as a measure of the effect size. The level of significance of post-hoc comparisons was set at *p <* 0.05. All statistical analyses were performed using custom Python 3.8 scripts and the Pingouin package [68].

### Modeling of the uncertain wrist reaching experiment

We modeled the drift term **f** by the following system:

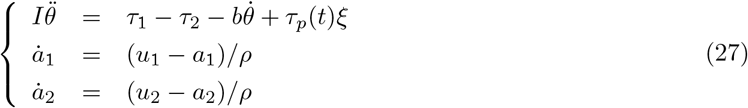

where *u*_1_, *u*_2_ ∈ [0, 1] are the muscle inputs of the biceps and triceps respectively, *a*_1_, *a*_2_ are the corresponding muscle activations, *τ*_1_, *τ*_2_ are the corresponding muscle torques and 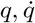 are the joint angle/velocity. The parameters *I* = 0.01 kg.m2 and *b* = 0.05 Nm.s/rad are respectively the moment of inertia about the elbow and the joint’s viscosity, and *ρ* = 0.04 s is the response time for muscle activations. As before, *ξ* was a Bernoulli variable with probability *α* and *τ*_*p*_ was defined as in Eq. 24 with *τ*_*p*_(*t*) = *τ*_*e*_(*t* − 0.8*T*) where *T* is the movement duration. This value was taken from experimental data which showed that the disturbance was activated around 80% of the total duration on average. The muscle torques were defined as in [3] to capture their variable viscoelasticity:

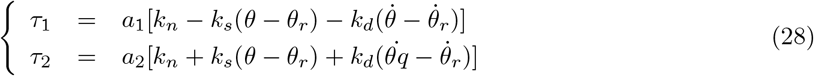

In the above equation, 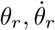 were taken from a reference trajectory built from a minimum jerk solution [69] and we set *k*_*n*_ = 15 Nm, *k*_*s*_ = 15 Nm/rad and *k*_*d*_ = 1.5 Nm/rad/s based on order of magnitudes found in the literature [31, 70, 71].

By denoting the state as 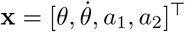 and the control as **u** = [*u*_1_, *u*_2_]^⊤^, the system of Eq. 27 can be written as a drift **f** (**x, u**) in the form of Eq. 1.

The diffusion term was modeled by the matrix:

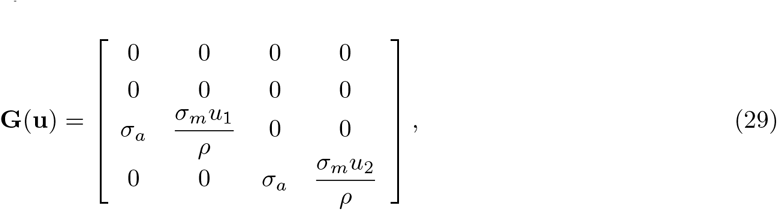

where *σ*_*a*_ = 0.02% and *σ*_*m*_ = 2% represent the magnitudes of additive noise and multiplicative respectively. These values were chosen such that the predicted co-contraction in the 0% condition and the rate of success overall matched the experimental values.

The task was to move the wrist from an initial random state **x**(0) = **x**_0_ ∼ *𝒩* (**m**_0_, **P**_0_) with **m**_0_ = [−*π/*6, 0, 10^−4^, 10^−4^]^⊤^ and **P**_0_ = 10^−5^**I** to **m**_*T*_ = [*π/*6, 0, *·, ·*]^⊤^ where *·* stands for undefined/free values.

The objective was to minimize the expectation of a quadratic expected cost mixing error and effort terms as follows:

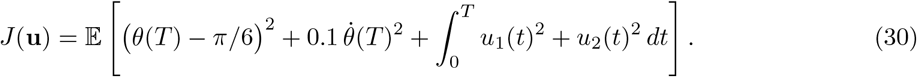

In all cases, the associated deterministic optimal control problems were solved numerically using Julia 1.10, JuMP [72] and Ipopt [73]. More precisely, we transcribed the continuous problem into an NLP problem using a trapezoidal scheme [74]. The Cholesky decomposition was used to rewrite the covariance differential equation as in [75] so that simple box constraints on the augmented state could be used to ensure the positive definiteness of the covariance matrix along the trajectory. All figures were made with Makie [76]. The code used to generate all simulations and all figures is open-source [link to be added upon acceptance].

## References

1. Latash ML. Muscle coactivation: definitions, mechanisms, and functions. J Neurophysiol. 2018;120:88–104. doi:10.1152/jn.00084.2018.

2. Voloshina AS, Kuo AD, Daley MA, Ferris DP. Biomechanics and energetics of walking on uneven terrain. Journal of Experimental Biology. 2013;doi:10.1242/jeb.081711.

3. Hogan N. Adaptive control of mechanical impedance by coactivation of antagonist muscles. IEEE Trans Autom Control. 1984;29(8):681–690.

4. Burdet E, Osu R, Franklin DW, Milner TE, Kawato M. The central nervous system sta- bilizes unstable dynamics by learning optimal impedance. Nature. 2001;414(6862):446–449. doi:10.1038/35106566.

5. Milner TE. Adaptation to destabilizing dynamics by means of muscle cocontraction. Exp Brain Res. 2002;143(4):406–416. doi:10.1007/s00221-002-1001-4.

6. Franklin DW, Liaw G, Milner TE, Osu R, Burdet E, Kawato M. Endpoint stiffness of the arm is directionally tuned to instability in the environment. J Neurosci. 2007;27(29):7705–7716.

7. Saliba CM, Rainbow MJ, Selbie WS, Deluzio KJ, Scott SH. Co-contraction uses dual control of agonist-antagonist muscles to improve motor performance. BioRxiv. 2020;doi:10.1101/2020.03.16.993527.

8. Börner H, Carboni G, Cheng X, Takagi A, Hirche S, Endo S, et al. Physically interacting hu- mans regulate muscle coactivation to improve visuo-haptic perception. Journal of Neurophysiology. 2023;129(2):494–499. doi:10.1152/jn.00420.2022.

9. Pruszynski JA, Kurtzer I, Lillicrap TP, Scott SH. Temporal evolution of “automatic gain-scaling”. J Neurophysiol. 2009;102(2):992–1003.

10. Crevecoeur F, Scott SH. Beyond Muscles Stiffness: Importance of State-Estimation to Account for Very Fast Motor Corrections. PLoS Computational Biology. 2014;10(10):e1003869. doi:10.1371/journal.pcbi.1003869.

11. Martino G, Beck ON, Ting LH. Voluntary muscle coactivation in quiet standing elicits reciprocal rather than coactive agonist-antagonist control of reactive balance. Journal of Neurophysiology. 2023;129(6):1378–1388. doi:10.1152/jn.00458.2022.

12. Gribble PL, Mullin LI, Cothros N, Mattar A. Role of cocontraction in arm movement accuracy. J Neurophysiol. 2003;89(5):2396–2405. doi:10.1152/jn.01020.2002.

13. Missenard O, Fernandez L. Moving faster while preserving accuracy. Neuroscience. 2011;197:233–241. doi:10.1016/j.neuroscience.2011.09.020.

14. Calalo JA, Roth AM, Lokesh R, Sullivan SR, Wong JD, Semrau JA, et al. The sensorimotor system modulates muscular co-contraction relative to visuomotor feedback responses to regulate movement variability. Journal of Neurophysiology. 2023;129(4):751–766. doi:10.1152/jn.00472.2022.

15. Todorov E, Jordan MI. Optimal feedback control as a theory of motor coordination. Nat Neurosci. 2002;5(11):1226–1235. doi:10.1038/nn963.

16. Li W, Todorov E. Iterative linearization methods for approximately optimal control and estimation of non-linear stochastic system. Int J Control. 2007;80(9):1439–1453.

17. van Beers RJ, Haggard P, Wolpert DM. The role of execution noise in movement variability. J Neurophysiol. 2004;91(2):1050–1063. doi:10.1152/jn.00652.2003.

18. Faisal AA, Selen LPJ, Wolpert DM. Noise in the nervous system. Nat Rev Neurosci. 2008;9(4):292–303. doi:10.1038/nrn2258.

19. Franklin DW, Burdet E, Tee KP, Osu R, Chew CM, Milner TE, et al. CNS learns stable, accurate, and efficient movements using a simple algorithm. Journal of neuroscience. 2008;28(44):11165– 11173.

20. Berret B, Jean F. Efficient computation of optimal open-loop controls for stochastic systems. Automatica. 2020;115:108874. 10.1016/j.automatica.2020.108874.

21. Berret B, Jean F. Stochastic optimal open-loop control as a theory of force and impedance planning via muscle co-contraction. PLOS Computational Biology. 2020;16(2):e1007414. doi:10.1371/journal.pcbi.1007414.

22. Koelewijn AD, Van Den Bogert AJ. Antagonistic co-contraction can minimize muscular effort in systems with uncertainty. PeerJ. 2022;10:e13085. doi:10.7717/peerj.13085.

23. Van Wouwe T, Ting LH, De Groote F. An approximate stochastic optimal control framework to simulate nonlinear neuro-musculoskeletal models in the presence of noise. PLOS Computational Biology. 2022;18(6):e1009338. doi:10.1371/journal.pcbi.1009338.

24. Takahashi CD, Scheidt RA, Reinkensmeyer DJ. Impedance Control and Internal Model Formation When Reaching in a Randomly Varying Dynamical Environment. Journal of Neurophysiology. 2001;86(2):1047–1051. doi:10.1152/jn.2001.86.2.1047.

25. Davidson PR, Wolpert DM. Motor learning and prediction in a variable environment. Current Opinion in Neurobiology. 2003;13(2):232–237. doi:10.1016/s0959-4388(03)00038-2.

26. Wei K, Wert D, Körding K. The Nervous System Uses Nonspecific Motor Learning in Response to Random Perturbations of Varying Nature. Journal of Neurophysiology. 2010;104(6):3053–3063. doi:10.1152/jn.01025.2009.

27. Ganesh G, Haruno M, Kawato M, Burdet E. Motor Memory and Local Minimization of Error and Effort, Not Global Optimization, Determine Motor Behavior. Journal of Neurophysiology. 2010;104(1):382–390. doi:10.1152/jn.01058.2009.

28. Hadjiosif AM, Smith MA. Flexible Control of Safety Margins for Action Based on Environmental Variability. Journal of Neuroscience. 2015;35(24):9106–9121. doi:10.1523/jneurosci.1883-14.2015.

29. Franklin DW, Osu R, Burdet E, Kawato M, Milner TE. Adaptation to stable and unstable dynamics achieved by combined impedance control and inverse dynamics model. J Neurophysiol. 2003;90:3270–3282. doi:10.1152/jn.01112.2002.

30. Crevecoeur F, Scott SH, Cluff T. Robust Control in Human Reaching Movements: A ModelFree Strategy to Compensate for Unpredictable Disturbances. The Journal of Neuroscience. 2019;39(41):8135–8148. doi:10.1523/jneurosci.0770-19.2019.

31. Bennett DJ, Hollerbach JM, Xu Y, Hunter IW. Time-varying stiffness of human elbow joint during cyclic voluntary movement. Exp Brain Res. 1992;88:433–442.

32. Berret B, Conessa A, Schweighofer N, Burdet E. Stochastic optimal feedforward-feedback control determines timing and variability of arm movements with or without vision. PLOS Computational Biology. 2021;17(6):e1009047. doi:10.1371/journal.pcbi.1009047.

33. Oksendal B. Stochastic Differential Equations. 4th ed. Springer Berlin; 1995.

34. Harris CM, Wolpert DM. Signal-dependent noise determines motor planning. Nature. 1998;394(6695):780–784. doi:10.1038/29528.

35. Todorov E. Stochastic optimal control and estimation methods adapted to the noise characteristics of the sensorimotor system. Neural Comput. 2005;17(5):1084–1108. doi:10.1162/0899766053491887.

36. Ruths J, Li JS. Optimal Control of Inhomogeneous Ensembles. IEEE Transactions on Automatic Control. 2012;57(8):2021–2032. doi:10.1109/tac.2012.2195920.

37. Phelps C, Gong Q, Royset JO, Walton C, Kaminer I. Consistent approximation of a nonlinear optimal control problem with uncertain parameters. Automatica. 2014;50(12):2987–2997. doi:10.1016/j.automatica.2014.10.025.

38. Ross IM, Proulx RJ, Karpenko M, Gong Q. Riemann–Stieltjes Optimal Control Problems for Uncertain Dynamic Systems. Journal of Guidance, Control, and Dynamics. 2015;38(7):1251–1263. doi:10.2514/1.g000505.

39. Phelps C, Royset JO, Gong Q. Optimal Control of Uncertain Systems Using Sample Average Approximations. SIAM Journal on Control and Optimization. 2016;54(1):1–29. doi:10.1137/140983161.

40. Shaffer R, Karpenko M, Gong Q. Relationships Between Unscented and Sensitivity Function-Based Optimal Control for Space Flight; 2017.

41. Walton C, Kaminer I, Gong Q. Consistent numerical methods for state and control constrained trajectory optimisation with parameter dependency. International Journal of Control. 2020;94(9):2564–2574. doi:10.1080/00207179.2020.1717633.

42. Katayama M, Kawato M. Virtual trajectory and stiffness ellipse during multijoint arm movement predicted by neural inverse models. Biol Cybern. 1993;69:353–362.

43. Corcos DM, Gottlieb GL, Agarwal GC. Organizing principles for single-joint movements. II. A speed-sensitive strategy. Journal of Neurophysiology. 1989;62(2):358–368. doi:10.1152/jn.1989.62.2.358.

44. Diedrichsen J, Shadmehr R, Ivry RB. The coordination of movement: optimal feedback control and beyond. Trends Cogn Sci. 2009;doi:10.1016/j.tics.2009.11.004.

45. Scott SH. The computational and neural basis of voluntary motor control and planning. Trends in cognitive sciences. 2012;16:541–549. doi:10.1016/j.tics.2012.09.008.

46. Ueyama Y. Mini-max feedback control as a computational theory of sensorimotor control in the presence of structural uncertainty. Front Comput Neurosci. 2014;8:119.

47. Kühn J, Bagnato C, Burdet E, Haddadin S. Arm movement adaptation to concurrent pain constraints. Scientific Reports. 2021;11(1). doi:10.1038/s41598-021-86173-7.

48. Shadmehr R, Wise SP. The Computational Neurobiology of Reaching and Pointing: A Foundation for Motor Learning. MIT press; 2005.

49. Wolpert DM, Flanagan JR. Motor learning. Current Biology. 2010;20(11):R467–R472. doi:10.1016/j.cub.2010.04.035.

50. Körding KP, Wolpert DM. Bayesian integration in sensorimotor learning. Nature. 2004;427(6971):244–247. doi:10.1038/nature02169.

51. Franklin DW, Wolpert DM. Computational mechanisms of sensorimotor control. Neuron. 2011;72(3):425–442.

52. Franklin DW, So U, Burdet E, Kawato M. Visual feedback is not necessary for the learning of novel dynamics. PLoS One. 2007;2(12):e1336.

53. Athans M. The Role and Use of the Stochastic Linear-Quadratic-Gaussian Problem in Control System Design. IEEE Trans Autom Control. 1971;16(6):529–552.

54. Mayne DQ, Rawlings JB, Rao CV, Scokaert PO. Constrained model predictive control: Stability and optimality. Automatica. 2000;36(6):789–814.

55. Mehrabi N, Sharif Razavian R, Ghannadi B, McPhee J. Predictive Simulation of Reaching Moving Targets Using Nonlinear Model Predictive Control. Frontiers in Computational Neuroscience. 2017;10. doi:10.3389/fncom.2016.00143.

56. Takagi A, Gomi H, Burdet E, Koike Y. A model predictive control strategy to regulate movements and interactions. BioRxiv. 2022;doi:10.1101/2022.08.24.505193.

57. Guigon E. A computational theory for the production of limb movements. Psychological Review. 2023;130(1):23–51. doi:10.1037/rev0000323.

58. Wolpert DM, Ghahramani Z, Jordan MI. An internal model for sensorimotor integration. Science. 1995;269(5232):1880–1882.

59. Kawato M. Internal models for motor control and trajectory planning. Curr Opin Neurobiol. 1999;9(6):718–727.

60. Wolpert DM, Ghahramani Z. Computational principles of movement neuroscience. Nat Neurosci. 2000;3 Suppl:1212–1217. doi:10.1038/81497.

61. Tauber F, Desmulliez M, Piccin O, Stokes AA. Perspective for soft robotics: the field’s past and future. Bioinspiration & Biomimetics. 2023;18(3):035001. doi:10.1088/1748-3190/acbb48.

62. Vanderborght B, Albu-Schaeffer A, Bicchi A, Burdet E, Caldwell D, Carloni R, et al. Variable impedance actuators: Moving the robots of tomorrow. In: Proc. IEEE/RSJ Int. Conf. Intelligent Robots and Systems; 2012. p. 5454–5455.

63. Li Y, Sena A, Wang Z, Xing X, Babič J, van Asseldonk E, et al. A review on interaction control for contact robots through intent detection. Progress in Biomedical Engineering. 2022;4(3):032004. doi:10.1088/2516-1091/ac8193.

64. Särkkä S, Solin A. Applied Stochastic Differential Equations. Institute of Mathematical Statistics Textbooks. Cambridge University Press; 2019.

65. Leparoux C, Bonalli R, Hérissé B, Jean F. Statistical linearization for robust motion planning. Systems & Control Letters. 2024;189:105825. 10.1016/j.sysconle.2024.105825.

66. Hermens HJ, Research R, Development BV, editors. European recommendations for surface ElectroMyoGraphy: results of the SENIAM project. No. 8 in SENIAM. Roessingh Research and Development; 1999.

67. Potvin JR, Brown SHM. Less is more: high pass filtering, to remove up to 99% of the surface EMG signal power, improves EMG-based biceps brachii muscle force estimates. Journal of Electromyography and Kinesiology. 2004;14(3):389–399. doi:10.1016/j.jelekin.2003.10.005.

68. Vallat R. Pingouin: statistics in Python. Journal of Open Source Software. 2018;3(31):1026. doi:10.21105/joss.01026.

69. Hogan N. An organizing principle for a class of voluntary movements. J Neurosci. 1984;4(11):2745– 2754.

70. Milner T, Cloutier C, Leger A, Franklin D. Inability to activate muscles maximally during cocontraction and the effect on joint stiffness. Experimental Brain Research. 1995;107(2). doi:10.1007/bf00230049.

71. Yoshii Y, Yuine H, Kazuki O, Tung Wl, Ishii T. Measurement of wrist flexion and extension torques in different forearm positions. BioMedical Engineering OnLine. 2015;14(1). doi:10.1186/s12938-015-0110-9.

72. Lubin M, Dowson O, Dias Garcia J, Huchette J, Legat B, Vielma JP. JuMP 1.0: Recent improvements to a modeling language for mathematical optimization. Mathematical Programming Computation. 2023;doi:10.1007/s12532-023-00239-3.

73. Wächter A, Biegler LT. On the implementation of an interior-point filter line-search algorithm for large-scale nonlinear programming. Mathematical Programming. 2005;106(1):25–57. doi:10.1007/s10107-004-0559-y.

74. Betts JT. Practical Methods for Optimal Control and Estimation Using Nonlinear Programming, Second Edition. 2nd ed. Society for Industrial and Applied Mathematics; 2010.

75. Särkkä S. On Unscented Kalman Filtering for State Estimation of Continuous-Time Nonlinear Systems. IEEE Transactions on Automatic Control. 2007;52(9):1631–1641. doi:10.1109/tac.2007.904453.

76. Danisch S, Krumbiegel J. Makie.jl: Flexible high-performance data visualization for Julia. Journal of Open Source Software. 2021;6(65):3349. doi:10.21105/joss.03349.

