## Supplementary Information Text for "Co-Contraction Embodies Uncertainty: An Optimal Feedforward Strategy for Robust Motor Control"

In this Supporting Information Text, we first give details about the case of a discrete random variable which introduces an alternative method of resolution. We then provide the proofs related to the analytical toy example and Proposition 1 of the main text.

#### Case of a discrete random variable $\boldsymbol{\xi}$

When $\boldsymbol{\xi}$ is a discrete random variable, we can address the problem differently. Let us assume that the random variable can take the values $\boldsymbol{\xi}_i$ for $i = 1, ..., s$ with probability $\alpha_i$, and we denote $\mathbf{x}^{\xi_i}(\cdot)$ the $n$-dimensional trajectory associated with the control $\mathbf{u}(\cdot)$ and the value $\boldsymbol{\xi}_i$. Given the nature of the problem, the expected cost in Eq. 2 of the main text can be rewritten

$$J(\mathbf{u}) = \sum_{i=1}^{s} \alpha_i q_f(\mathbf{x}^{\xi_i}(T), T) + \int_0^T \sum_{i=1}^{s} \alpha_i q(\mathbf{x}^{\xi_i}(t), \mathbf{u}(t), t)\, dt, \tag{1}$$

where $q_f$ and $q$ are quadratic functions in the state.

If we augment the state of the system by setting $\tilde{\mathbf{x}} = (\mathbf{x}^{\xi_1}, \ldots, \mathbf{x}^{\xi_s})$, the cost can be written more compactly as

$$J(\mathbf{u}) = \tilde{q}_f(\tilde{\mathbf{x}}(T), T) + \int_0^T \tilde{q}(\tilde{\mathbf{x}}(t), \mathbf{u}(t), t)\, dt, \tag{2}$$

where the function $\tilde{q}(\tilde{\mathbf{x}}(t), \mathbf{u}(t), t) = \sum_{i=1}^{s} \alpha_i q(\mathbf{x}^{\xi_i}(t), \mathbf{u}(t), t)$ and $\tilde{q}_f(\tilde{\mathbf{x}}(T), T) = \sum_{i=1}^{s} \alpha_i q_f(\mathbf{x}^{\xi_i}(T), T)$.

The initial uncertain optimal control problem with discrete random uncertainty $\boldsymbol{\xi}$ is equivalent to a deterministic optimal control problem in the augmented state $\tilde{\mathbf{x}} \in \mathbb{R}^{ns}$ with a dynamical system of the form

$$\dot{\tilde{\mathbf{x}}}(t) = \tilde{\mathbf{f}}(\tilde{\mathbf{x}}(t), \mathbf{u}(t), t) \tag{3}$$

where $\tilde{\mathbf{f}} = (\mathbf{f}(\mathbf{x}^{\xi_1}(t), \mathbf{u}(t), t; \boldsymbol{\xi}_1), \ldots, \mathbf{f}(\mathbf{x}^{\xi_s}(t), \mathbf{u}(t), t; \boldsymbol{\xi}_s))$.

*Remark.* There is no restriction on the type of distribution for $\boldsymbol{\xi}$ and on the type of dynamics in this approach. The main limitation is the size of the augmented space ($ns$), which may limit the relevance of the approach when $s$ is too large. In this case, the approach proposed in Problem 3 of the main text may be more suited. If the system is governed by the nonlinear SDE

$$d\mathbf{x}_t = \mathbf{f}(\mathbf{x}_t, \mathbf{u}(t), t; \boldsymbol{\xi})\, dt + \mathbf{G}(\mathbf{x}_t, \mathbf{u}(t), t)\, d\mathbf{w}_t$$

where $\boldsymbol{\xi}$ a discrete random variable, the above approach can be combined with the SOOC approach of [1,2], that is, we can introduce the variables $\mathbf{m}^{\boldsymbol{\xi}_i}(t)$ and $\mathbf{P}^{\boldsymbol{\xi}_i}(t)$ for $i = 1..s$.

#### Proof of the toy stabilization task

To illustrate that random external disturbances lead to impedance control, let us consider a toy stabilization task where analytical computations are tractable. We consider the bilinear system

$$\dot{x}(t) = f(t) - k(t)x(t) + \xi, \tag{4}$$

where $x$ is the scalar state, $\mathbf{u} = [f, k]^\top$ is the control vector composed of a term $f$ representing a force and a term $k$ representing a stiffness, and $\xi$ is a random external disturbance.

Here the random variable $\xi$ corresponds to an external force that can be applied or not, so that $\xi$ equals 0 with probability $\alpha$ and 1 with probability $1 - \alpha$.

We assume as previously that the control law $\mathbf{u}(t)$ is open-loop, and is determined as the one that minimizes the expected cost with scalar weights $q \geq 1$, $q_f > 0$,

$$J(\mathbf{u}) = \mathbb{E}\left[q_f x(T)^2 + \int_0^T \left(f(t)^2 + k(t)^2 + qx(t)^2\right)\, dt\right],$$

among the *open-loop* controls $\mathbf{u}(t)$ ensuring that the expectation $\mathbb{E}_\xi[x(T)]$ of the final state equals 0. We assume that the initial state is $x(0) = 0$ and that time $T$ is fixed.

We will now prove that the very presence of uncertainty leads to impedance control in this problem.

*First case (without uncertainty).* Consider the cases where there is no uncertainty, that is when $\alpha = 0$ or $\alpha = 1$. It consists in minimizing the cost

$$J(\mathbf{u}) = q_f x(T)^2 + \int_0^T \left(f(t)^2 + k(t)^2 + qx(t)^2\right)\, dt,$$

among the trajectories $(x, \mathbf{u})$ of the deterministic controlled system

$$\dot{x}(t) = f(t) - k(t)x(t) + \alpha,$$

satisfying $x(0) = x(T) = 0$.

Let us apply the necessary conditions of the Pontryagin's Maximum Principle (PMP) (see [3]). The Hamiltonian function is

$$\mathcal{H}(x, \lambda, \mathbf{u}) = \lambda(f - kx + \alpha) - \frac{1}{2}\left(f^2 + k^2 + qx^2\right).$$

On the one hand, each value of the optimal control $\mathbf{u}$ must maximize $\mathcal{H}$ with respect to the control, which yields $f = \lambda, k = -\lambda x$. On the other hand, the optimal trajectory and its coadjoint variable $\lambda$ must satisfy the Hamiltonian differential equations

$$\dot{x} = \lambda(1 + x^2) + \alpha, \qquad \dot{\lambda} = x(q - \lambda^2).$$

Taking the derivative of $\dot{x}$, we obtain

$$\ddot{x} = \begin{cases} x(1 + x^2)(q + \lambda^2) & \text{if } \alpha = 0, \\ x\left(x^2(q + \lambda^2) + (1 + \lambda)^2 + q - 1\right) & \text{if } \alpha = 1. \end{cases}$$

Thus $x\ddot{x} \geq 0$ (recall $q \geq 1$) which implies that the function $t \mapsto (x\dot{x})(t)$ is nondecreasing. Since this function is zero at $t = 0$, it is nonnegative, therefore $x^2(t)$ is nondecreasing. The condition $x(0) = x(T) = 0$ then implies that the optimal trajectory is $x \equiv 0$ and the optimal control satisfies $f \equiv -\alpha, k \equiv 0$, showing that there is no use of stiffness (or impedance control) in these cases.

*Second case (with uncertainty)*. Let us now show that the above situation ($k \equiv 0$) never appears in the presence of uncertainty, that is, when $0 < \alpha < 1$.

We first convert the optimal control problem into a deterministic one by augmenting the state of the system. We set $\mathbf{x} = (x_0, x_1)$, with an augmented dynamics

$$\dot{x}_0(t) = f(t) - k(t)x_0(t), \qquad \dot{x}_1(t) = f(t) - k(t)x_1(t) + 1, \tag{5}$$

initial conditions $\mathbf{x}(0) = 0$, and a terminal condition $\mathbb{E}_\xi[x(T)] = (1 - \alpha)x_0(T) + \alpha x_1(T) = 0$. The cost writes as

$$J(\mathbf{u}) = q_f((1 - \alpha)x_0(T)^2 + \alpha x_1(T)^2) + \int_0^T \left(f(t)^2 + k(t)^2 + q(1 - p)x_0(t)^2 + qpx_1(t)^2\right) dt.$$

The solutions of the corresponding optimal control problem must satisfy the necessary condition given by the PMP. Define the Hamiltonian function

$$\mathcal{H}(\mathbf{x}, \boldsymbol{\lambda}, \mathbf{u}, \nu) = \lambda_0(f - kx_0) + \lambda_1(f - kx_1 + 1) - \nu\left(f^2 + k^2 + q(1 - \alpha)x_0^2 + q\alpha x_1^2\right). \tag{6}$$

If $\mathbf{u}(t)$ is an optimal control with associated trajectory $\mathbf{x}(t)$, then there exist $\nu = 0$ or $1/2$ and a function

$\boldsymbol{\lambda}(t) = (\lambda_0(t), \lambda_1(t))$ such that $(\nu, \boldsymbol{\lambda}) \neq 0$ and, for every $t \in [0, T]$,

$$\dot{\lambda}_0 = -\frac{\partial \mathcal{H}}{\partial x_0}(\mathbf{x}, \boldsymbol{\lambda}, \mathbf{u}, \nu), \quad \dot{\lambda}_1 = -\frac{\partial \mathcal{H}}{\partial x_1}(\mathbf{x}, \boldsymbol{\lambda}, \mathbf{u}, \nu), \tag{7}$$

$$\mathcal{H}(\mathbf{x}(t), \boldsymbol{\lambda}(t), \mathbf{u}(t), \nu) = \max_{\mathbf{v}} \mathcal{H}(\mathbf{x}(t), \boldsymbol{\lambda}(t), \mathbf{v}, \nu), \tag{8}$$

plus conditions of transversality that we do not need here. A simple computation shows that $\nu = 0$ leads to a contradiction, so $\nu = 1/2$ and the maximization condition on $\mathcal{H}$ is equivalent to

$$f = \lambda_0 + \lambda_1, \qquad k = -\lambda_0 x_0 - \lambda_1 x_1, \tag{9}$$

whereas the Hamiltonian differential equations write as

$$\dot{\lambda}_0 = q(1-\alpha)x_0 + k\lambda_0, \qquad \dot{\lambda}_1 = q\alpha x_1 + k\lambda_1. \tag{10}$$

Assume by contradiction that there exists an optimal solution with no stiffness, that is, an optimal control with $k \equiv 0$. This implies by Eqs. 5, 9 and 10 that there exist a solution $(\mathbf{x}(t), \boldsymbol{\lambda}(t))$ of

$$\begin{cases} \dot{x}_0 = \lambda_0 + \lambda_1, \\ \dot{x}_1 = \lambda_0 + \lambda_1 + 1, \end{cases} \quad x_0(0) = x_1(0) = 0, \quad \begin{cases} \dot{\lambda}_0 = q(1-\alpha)x_0, \\ \dot{\lambda}_1 = q\alpha x_1, \end{cases} \tag{11}$$

verifying $k = -\lambda_0 x_0 - \lambda_1 x_1 \equiv 0$. A simple computation shows that the solutions of the above differential equations are of the form

$$\begin{cases} x_0 = -\alpha t + a \sinh(\sqrt{q}t), \\ x_1 = (1-\alpha)t + a\sinh(\sqrt{q}t), \end{cases} \qquad \begin{cases} \lambda_0 = b_0 - q\frac{\alpha(1-\alpha)}{2}t^2 + a\sqrt{q}(1-\alpha)\cosh(\sqrt{q}t), \\ \lambda_1 = b_1 + q\frac{\alpha(1-\alpha)}{2}t^2 + a\sqrt{q}\alpha\cosh(\sqrt{q}t), \end{cases} \tag{12}$$

for some constants $a, b_0, b_1$. As a consequence, $k = -\lambda_0 x_0 - \lambda_1 x_1$ writes as

$$k = (\alpha b_0 - (1-\alpha)b_1)\, t - q\frac{\alpha(1-\alpha)}{2}t^3 + a(b_0+b_1)\sinh(\sqrt{q}t) - a^2\sqrt{q}\sinh(\sqrt{q}t)\cosh(\sqrt{q}t), \tag{13}$$

and cannot be identically zero when $0 < \alpha < 1$, whatever the values of the constants $a, b_0, b_1$.

We thus get the conclusion that uncertainty ($0 < \alpha < 1$) leads to some nonzero impedance control ($k \not\equiv 0$). Further note that the random variables has mean $\alpha$ and variance $\alpha(1-\alpha)$ so that it can be seen that the level of stiffness directly depends on the variance (that is, the degree of task uncertainty).

#### Proof of Proposition 1 of the main text

The Proposition results from the two following lemmas.

**Lemma 1.** *Assume that the dynamics* $\mathbf{f}$ *is smooth with compact support and depends affinely of the* *random parameter* $\boldsymbol{\xi}$*, i.e.*

$$\mathbf{f}(\mathbf{x},\mathbf{u};\boldsymbol{\xi}) = \mathbf{g}(\mathbf{x},\mathbf{u}) + \mathbf{G}(\mathbf{u})\boldsymbol{\xi}.$$

*Then there exists a constant* $C > 0$ *such that, for any* $t \in [0,T]$*,*

$$\sup_{s\in[0,t]} \|\mathbf{m}_{\mathbf{x}}(s) - \mathbf{m}(s)\|^2 + \sup_{s\in[0,t]} \|\mathbf{P}_{\mathbf{x}}(s) - \mathbf{P}(s)\| \leq C \left( \sup_{s\in[0,t]} \|\mathbf{P}(s)\| + \sup_{s\in[0,t]} \|\mathbf{D}(s)\| \right).$$

*Proof.* Note first that, by a direct computation, $\mathbf{m}_{\mathbf{x}} = \mathbb{E}[\mathbf{x}]$, $\mathbf{P}_{\mathbf{x}} = \mathbb{E}[(\mathbf{x} - \mathbf{m}_{\mathbf{x}})(\mathbf{x} - \mathbf{m}_{\mathbf{x}})^\top]$ and $\mathbf{D}_{\mathbf{x}} = \mathbb{E}[(\mathbf{x} - \mathbf{m}_{\mathbf{x}})(\boldsymbol{\xi} - \boldsymbol{\mu})^\top]$ satisfy

$$\begin{aligned}
\dot{\mathbf{m}}_{\mathbf{x}} = \quad & \mathbb{E}[\mathbf{g}(\mathbf{x},\mathbf{u})] + \mathbf{G}(\mathbf{u})\boldsymbol{\mu}, \\
\dot{\mathbf{D}}_{\mathbf{x}} = \quad & \mathbb{E}\left[ \big(\mathbf{g}(\mathbf{x},\mathbf{u}) - \mathbb{E}[\mathbf{g}(\mathbf{x},\mathbf{u})]\big)\big(\boldsymbol{\xi} - \boldsymbol{\mu}\big)^\top \right] + \mathbf{G}(\mathbf{u})\boldsymbol{\Sigma}, \\
\dot{\mathbf{P}}_{\mathbf{x}} = \quad & \mathbb{E}\left[ \big(\mathbf{g}(\mathbf{x},\mathbf{u}) - \mathbb{E}[\mathbf{g}(\mathbf{x},\mathbf{u})]\big)\big(\mathbf{x} - \mathbf{m}_{\mathbf{x}}\big)^\top \right] + \mathbb{E}\left[ \big(\mathbf{x} - \mathbf{m}_{\mathbf{x}}\big)\big(\mathbf{g}(\mathbf{x},\mathbf{u}) - \mathbb{E}[\mathbf{g}(\mathbf{x},\mathbf{u})]\big)^\top \right] \\
& + \mathbf{G}(\mathbf{u})\mathbf{D}_{\mathbf{x}}^\top + \mathbf{D}_{\mathbf{x}}\mathbf{G}(\mathbf{u})^\top.
\end{aligned} \tag{14}$$

whereas $(\mathbf{m},\mathbf{P},\mathbf{D})$ satisfies Eq. 14, and $(\mathbf{m}_{\mathbf{x}},\mathbf{P}_{\mathbf{x}},\mathbf{D}_{\mathbf{x}})(0) = (\mathbf{m},\mathbf{P},\mathbf{D})(0)$.

We will use a Taylor expansion with integral rest of the function $\mathbf{g}$,

$$\mathbf{g}(\mathbf{x},\mathbf{u}) = \mathbf{g}(\mathbf{m},\mathbf{u}) + \mathbf{h}(\mathbf{m},\mathbf{x},\mathbf{u})(\mathbf{x} - \mathbf{m}),$$

the function $\mathbf{h}$ being smooth with compact support. Thus,

$$\|\dot{\mathbf{m}}_{\mathbf{x}} - \dot{\mathbf{m}}\| \leq \|\mathbb{E}\left[\mathbf{g}(\mathbf{x},\mathbf{u})\right] - \mathbf{g}(\mathbf{m},\mathbf{u})\| \leq \|\mathbb{E}\left[\mathbf{h}(\mathbf{m},\mathbf{x},\mathbf{u})(\mathbf{x} - \mathbf{m})\right]\| \leq C\|\mathbb{E}\left[\mathbf{x} - \mathbf{m}\right]\|$$

This inequality, together with Cauchy-Schwarz and Jensen inequalities, yields to

$$\begin{aligned}
\sup_{s\in[0,t]} \|\mathbf{m}_{\mathbf{x}}(s) - \mathbf{m}(s)\|^2 &\leq \left( \int_0^t \|\dot{\mathbf{m}}_{\mathbf{x}}(s) - \dot{\mathbf{m}}(s)\|\, ds \right)^2 \\
&\leq C \left( \int_0^t \|\mathbb{E}\left[\mathbf{x}(s) - \mathbf{m}(s)\right]\|\, ds \right)^2 \\
&\leq C \int_0^t \|\mathbb{E}\left[\mathbf{x}(s) - \mathbf{m}(s)\right]^2\|\, ds.
\end{aligned}$$

Writing $\mathbf{x} - \mathbf{m}$ as $\mathbf{x} - \mathbf{m}_{\mathbf{x}} + \mathbf{m}_{\mathbf{x}} - \mathbf{m}$ and using the fact that $\mathrm{tr}(\mathbf{P}_{\mathbf{x}}) \leq C\|\mathbf{P}_{\mathbf{x}}\|$, we obtain

$$\begin{aligned}
\sup_{s\in[0,t]} \|\mathbf{m}_{\mathbf{x}}(s) - \mathbf{m}(s)\|^2 &\leq C \int_0^t \left( \|\mathbf{m}_{\mathbf{x}}(s) - \mathbf{m}(s)\|^2 + \mathrm{tr}(\mathbf{P}_{\mathbf{x}}(s)) \right) ds, \\
&\leq C \left( \int_0^t \sup_{[0,s]} \|\mathbf{m}_{\mathbf{x}} - \mathbf{m}\|^2 ds + \sup_{s\in[0,t]} \|\mathbf{P}_{\mathbf{x}}(s)\| \right).
\end{aligned}$$

Finally, using Gronwall's inequality and $\mathbf{P_x} = \mathbf{P_x} - \mathbf{P} + \mathbf{P}$, we obtain

$$\sup_{s\in[0,t]} \|\mathbf{m_x}(s) - \mathbf{m}(s)\|^2 \leq C \left( \sup_{s\in[0,t]} \|\mathbf{P_x}(s) - \mathbf{P}(s)\| + \sup_{s\in[0,t]} \|\mathbf{P}(s)\| \right).$$

A similar estimate shows

$$\begin{aligned} \sup_{s\in[0,t]} \|\mathbf{P_x}(s) - \mathbf{P}(s)\| \leq & \quad C \left( \sup_{s\in[0,t]} \|\mathbf{D_x}(s) - \mathbf{D}(s)\| + \sup_{s\in[0,t]} \|\mathbf{P}(s)\| + \sup_{s\in[0,t]} \|\mathbf{D}(s)\| \right), \\ \sup_{s\in[0,t]} \|\mathbf{D_x}(s) - \mathbf{D}(s)\| \leq & \quad C \sup_{s\in[0,t]} \|\mathbf{D}(s)\|. \end{aligned}$$

Combining the last three inequalities, we obtain the lemma. □

**Lemma 2.** *Assume that the dynamics* $\mathbf{f}$ *is smooth with compact support. Then there exists a constant* $C > 0$ *such that, for any* $t \in [0, T]$,

$$\sup_{s\in[0,t]} \|\mathbf{m_x}(s) - \mathbf{m}(s)\|^2 + \sup_{s\in[0,t]} \|\mathbf{P_x}(s) - \mathbf{P}(s)\| \leq C \left( \sup_{s\in[0,t]} \|\mathbf{P}(s)\| + \sup_{s\in[0,t]} \|\mathbf{D}(s)\| + \|\mathbf{\Sigma}\| \right).$$

*Proof.* Set $\widetilde{\mathbf{x}} = (\mathbf{x}, \xi)$, $\widetilde{\boldsymbol{\xi}} = \mathbf{0}$, $\widetilde{\mathbf{x}}_0 = (\mathbf{x}_0, \boldsymbol{\xi})$, and

$$\widetilde{\mathbf{f}}(\widetilde{\mathbf{x}}, \mathbf{u}; \widetilde{\boldsymbol{\xi}}) = \begin{pmatrix} \mathbf{f}(\mathbf{x}, \mathbf{u}; \boldsymbol{\xi}) \\ \mathbf{0} \end{pmatrix}.$$

With the notations of the Materials and Methods, we have for the extended system:

$$\widetilde{\mathbf{m}}_\mathbf{x} = (\mathbf{m_x}, \boldsymbol{\mu}), \quad \widetilde{\mathbf{m}} = (\mathbf{m}, \boldsymbol{\mu}), \quad \widetilde{\mathbf{D}}_\mathbf{x} = \widetilde{\mathbf{D}} = \mathbf{0}, \quad \widetilde{\mathbf{P}}_\mathbf{x} = \begin{pmatrix} \mathbf{P_x} & \mathbf{D_x} \\ \mathbf{D_x}^\top & \mathbf{\Sigma} \end{pmatrix}, \quad \widetilde{\mathbf{P}} = \begin{pmatrix} \mathbf{P} & \mathbf{D} \\ \mathbf{D}^\top & \mathbf{\Sigma} \end{pmatrix}.$$

Since the dynamics $\widetilde{\mathbf{f}}$ is not perturbed by a random parameter, it can be considered as affine with respect to this parameter. Thus Lemma 1 applies and we obtain the desired estimate. □

### References


1. Berret B, Jean F. Efficient computation of optimal open-loop controls for stochastic systems. Automatica. 2020;115:108874. doi:https://doi.org/10.1016/j.automatica.2020.108874.

2. Berret B, Jean F. Stochastic optimal open-loop control as a theory of force and impedance planning via muscle co-contraction. PLOS Computational Biology. 2020;16(2):e1007414. doi:10.1371/journal.pcbi.1007414.

3. Trélat E. Contrôle optimal : Théorie & applications. Vuibert, editor; 2008.
